# Preclinical evaluation of a COVID-19 vaccine candidate based on a recombinant RBD fusion heterodimer of SARS-CoV-2

**DOI:** 10.1101/2021.11.22.469117

**Authors:** Antonio Barreiro, Antoni Prenafeta, Gregori Bech-Sabat, Mercè Roca, Eva Perozo Mur, Ricard March, Luis González-González, Laia Madrenas, Júlia Corominas, Alex Fernández, Alexandra Moros, Manuel Cañete, Mercè Molas, Thais Pentinat-Pelegrin, Clara Panosa, Alberto Moreno, Ester Puigvert Molas, Eva Pol Vilarrassa, Jordi Palmada, Carme Garriga, Teresa Prat Cabañas, Javier Iglesias-Fernández, Júlia Vergara-Alert, Cristina Lorca-Oró, Núria Roca, Leira Fernández-Bastit, Jordi Rodon, Mònica Pérez, Joaquim Segalés, Edwards Pradenas, Silvia Marfil, Benjamin Trinité, Raquel Ortiz, Bonaventura Clotet, Julià Blanco, Jorge Díaz Pedroza, Rosa Ampudia Carrasco, Yaiza Rosales Salgado, Jordina Loubat-Casanovas, Sara Capdevila Larripa, Julia Garcia Prado, Jordi Barretina, Marta Sisteré-Oró, Paula Cebollada Rica, Andreas Meyerhans, Laura Ferrer

**Author notes:** These authors contributed equally (Antonio Barreiro, Antoni Prenafeta).

## Abstract

Current C0VID-19 vaccines have been associated with a decline in infection rates, prevention of severe disease and a decrease in mortality rates. However, SARS-CoV-2 variants are continuously evolving, and development of new accessible COVID-19 vaccines is essential to mitigate the pandemic. Here, we present data on preclinical studies in mice of a receptor-binding domain (RBD)-based recombinant protein vaccine (PHH-1V) consisting of an RBD fusion heterodimer comprising the B.1.351 and B.1.1.7 SARS-CoV-2 variants formulated in SQBA adjuvant, an oil-in-water emulsion. A prime-boost immunisation with PHH-1V in BALB/c and K18-hACE2 mice induced a CD4^+^ and CD8^+^ T cell response and RBD-binding antibodies with neutralising activity against several variants, and also showed a good tolerability profile. Significantly, RBD fusion heterodimer vaccination conferred 100% efficacy, preventing mortality in SARS-CoV-2 infected K18-hACE2 mice, but also reducing Beta, Delta and Omicron infection in lower respiratory airways. These findings demonstrate the feasibility of this recombinant vaccine strategy.

## 1. Introduction

In December 2019, severe acute respiratory syndrome coronavirus 2 (SARS-CoV-2) was identified as the etiological agent of the coronavirus disease 2019 (COVID-19). Soon afterwards, the scientific community and the pharmaceutical industry began focusing on the development of effective COVID-19 vaccines to mitigate the health emergency. Thanks to these efforts, several vaccines are currently available, and more than 12.9 billion doses have been administered worldwide (November 2022)^1^. The decline in new infection rates in many countries coincides with the introduction of vaccines. However, COVID-19 cases continue to emerge, probably due to the appearance and evolution of SARS-CoV-2 variants, the decline of immunological protection provided by the current vaccines, and especially the lack of homogenous distribution of COVID-19 vaccines, with only 22.5% of people in low-income countries having received at least one dose (December 2021)^2,3^. As the global outbreak continues, the pandemic is far from being over and it is not clear if the available vaccines will be sufficient to revert the situation. Thus, it is still critical to develop second-generation vaccines using different platforms that are effective against new variants and that could be further used as a booster, particularly to maintain or even enhance immunity against SARS-CoV-2^4,5^. Moreover, it is of relevance that these recent vaccines can be stored in refrigerated conditions, making them easier to distribute and avoiding more expensive and less available ultra-low temperature storage and transport conditions to ensure their global supply. Currently authorised vaccines, whether approved under emergency use or fully licensed, are based on viral vectors, inactivated viruses, nucleic acid-based vaccines, and protein subunit vaccines^6^.

SARS-CoV-2 is a betacoronavirus belonging to the subfamily *Coronovirinae*, within the family *Coronaviridae* and the order *Nidovirales*. The SARS-CoV-2 genome is a positive-sense single-stranded RNA (+ssRNA) molecule. The genome size ranges between 27 and 32 kbp, one of the largest known RNA viruses. The genomic structure of SARS-CoV-2 contains at least six open reading frames (ORFs), encoding for at least four structural proteins, namely: envelope or spike (S) glycoprotein; membrane (M) proteins, responsible for the shaping of the virions; envelope (E) proteins, responsible for the virions assembly and release; and nucleocapsid (N) proteins, involved in the RNA genome packaging^7^. The trimeric S glycoprotein of SARS-CoV-2 is the primary target of viral neutralising antibodies and has been the main protein candidate for vaccine development^8^. Consistent with SARS-CoV, angiotensin-converting enzyme 2 (ACE2) binding of the S protein allows cellular entry of SARS-CoV-2 viral particles^9^. This protein consists of 2 domains, S1 and S2, allowing the binding of the viral particles and cellular entry by fusing with the host cell membrane^10^. The receptor-binding domain (RBD) (Thr333-Gly526) is found in the S1 domain and it contains a highly immunogenic receptor-binding motif (RBM) that directly interacts with ACE2 and neutralising antibodies^11^. Therefore, most key mutations are found in the RBM, allowing the virus to adapt and escape the previously developed immunity^12,13^. To date, several SARS-CoV-2 VoCs with key mutations in the S protein have emerged: Alpha (B.1.1.7), Beta (B.1.351), Gamma (P.1), Delta (B.1.617.2) and Omicron (B.1.1.529)^14,15^.

The S protein is the primary target in vaccine development against betacoronaviruses due to its accessibility for immune recognition^16^. It has been reported that two proline substitutions in the original S protein sequence (S-2P) of MERS-CoV, SARS-CoV and HKU1 coronavirus maintain the antigenic conformation but retain S proteins in the prototypical prefusion conformation^17^. Thus, learning from these previous results, this S-2P design is used in the licensed mRNA-based vaccines Comirnaty^®^ (Pfizer-BioNTech)^18^ and Spikevax® (Moderna)^19^, which substitute the K986 and V987 residues for prolines from the original S protein variant. Likewise, the adenoviral vector-based vaccine Jcovden^®^ (Johnson & Johnson) also contains DNA-encoding for the S-2P protein of SARS-CoV-2^20^.

Adjuvanted protein-based subunit vaccines represent an important type of vaccine, yet their development has lagged compared to other platforms due to the need to optimise the manufacturing process for each protein antigen. The most advanced subunit vaccine programme against COVID-19 is the Novavax vaccine (NVX-CoV-2373, marketed as Nuvaxovid^®^), which is produced in insect cells in combination with a saponin-based adjuvant (Matrix-M)^21^. This vaccine consists of the stable full-length S protein in the antigenically optimal prefusion conformation. In addition, the Sanofi-GSK vaccine, known as VidPrevtyn Beta^®^, consists of soluble prefusion-stabilised S trimers from SARS-CoV-2 produced in insect cells combined with the AS03 adjuvant^22^. Both vaccines have been tested in human clinical trials and have recently received an EU marketing authorisation^21,23^. Notably, recombinant proteins are competitive vaccine candidates with an adequate safety profile, no risk of genome integration, no live components, and suitable for people with compromised immune systems^24^, showing high productivity yields and good stability profiles^24–26^.

Most cloned neutralising antibodies target the RBD in the S1 domain^27^, although there are additional immunogenic epitopes outside this domain^28^. It has been reported that more than 90% of neutralising antibodies isolated from convalescent patients target the RBD in one-third of cases^29^. Moreover, depleted sera and plasma samples from individuals vaccinated with a 250-μg dose of the mRNA-1273 vaccine showed that up to 99% of neutralising antibodies target the RBD, even though the antigen is based on the whole prefusion spike conformation^30^. Given that the RBD domain of the S protein directly interacts with the ACE2 receptor, RBD-targeting antibodies are not expected to cause antibody-dependent enhancement (ADE), unlike non-neutralising or sub-neutralising antibodies^31^, highlighting the importance of the RBD in the immune response against SARS-CoV-2 and emphasising its relevance as a powerful and efficient immunogen in vaccine design.

In view of the inherent particularities of the S protein, and especially the RBD domain, our team developed a vaccine-candidate platform based on this immunogen. From amongst the tested preliminary vaccine candidates, combined with one or several adjuvants, we finally proceeded with a protein-based subunit vaccine candidate, namely PHH-1V, consisting of a recombinant RBD fusion heterodimer of the B.1.351 and B.1.1.7 variants of SARS-CoV-2 expressed in Chinese hamster ovary (CHO) cells and formulated with the squalene-based adjuvant (SQBA). Specifically, the SQBA adjuvant is an oil-in-water emulsion comprising well-known components which are used as adjuvants in human medicine^32^. Hence, the main aims of this study were to assess the safety and efficacy of the PHH-1V vaccine in BALB/c and K18-hACE2-transgenic mice models, and to characterise the RBD fusion heterodimer antigen and its immunogenicity.

## 2. Results

### 2.1. Recombinant RBD fusion heterodimer expression and characterisation

The antigen of the PHH-1V vaccine candidate is a recombinant RBD fusion heterodimer based on the B.1.351 (Beta) and B.1.1.7 (Alpha) SARS-CoV-2 variants (**Figure 1A**). The N-terminal monomer contains the amino acid sequence of the SARS-CoV-2 RBD protein from the B.1.351 variant, whereas the C-terminal monomer contains the amino acid sequence of the SARS-CoV-2 RBD protein from the B.1.1.7 variant. The rationale behind this antigenic construct was based on maximisation of the affinity constant towards its target receptor, allowing the accommodation of each RBD variant bound to a single hACE monomer within the same or a different receptor. Three-dimensional structural models generated with AlphaFold2^33^ highlighted the coexistence of two different conformations of the PHH-1V construct. More specifically, one of the conformations is characterised by a protein-protein interaction between both RBD variants, whereas the other presents separated RBD domains stabilised by interactions of the N-/C-terminal regions (**Figure 2B**). RBD monomer binding towards individual human ACE2 (hACE2) units requires the preferential adoption of a separated RBD stabilised conformation, and thus construct generation followed this requirement.

**Figure 1.**
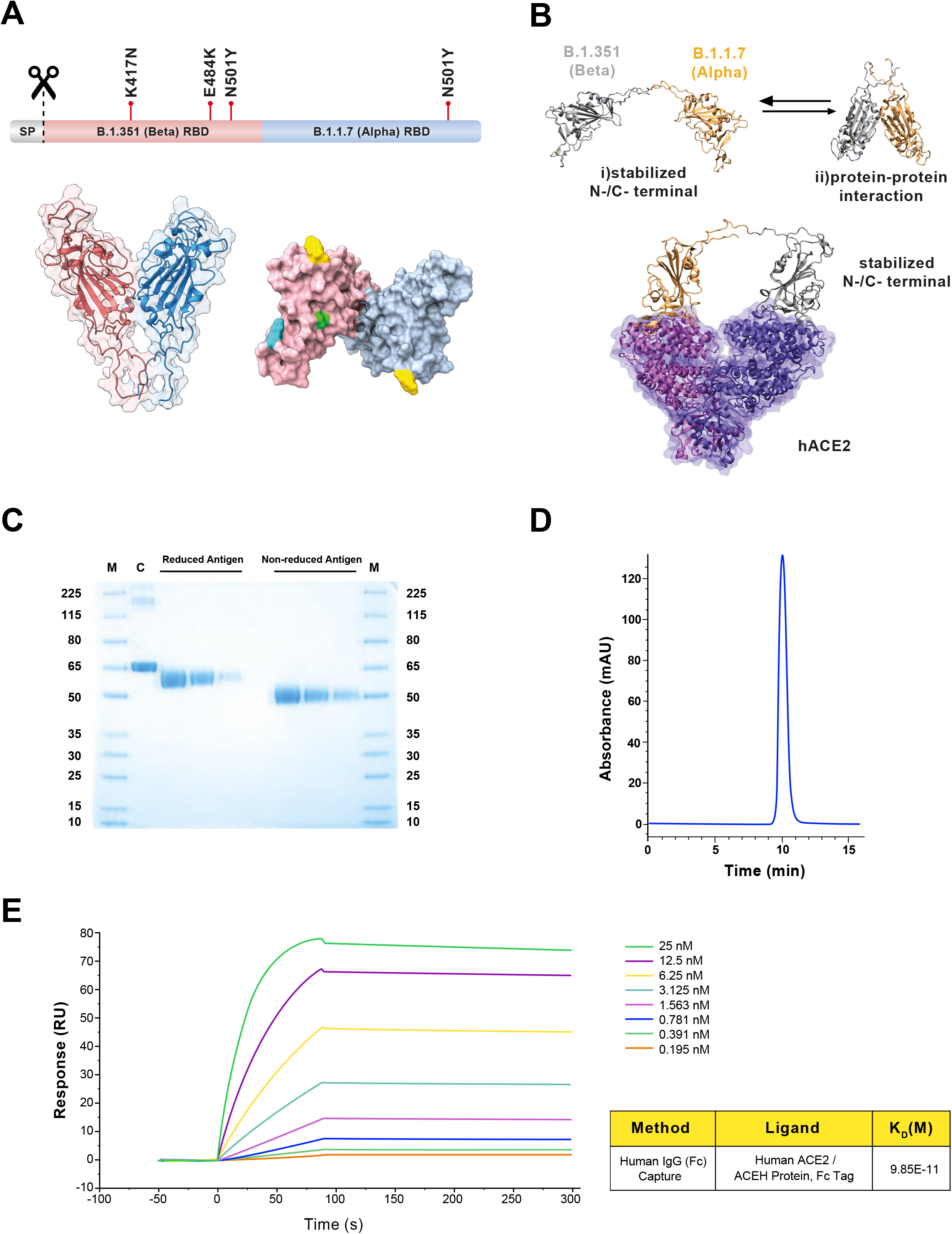
Structure and characterisation of the B.1.351 (Beta) - B.1.1.7 (Alpha) receptor-binding domain (RBD) heterodimer, immunogen of PHH-1V. (**A**) Structural representation of the RBD heterodimer. Top: sequence diagram. Bottom left: front view of the RBD heterodimer cartoon structure. Bottom right: top view of the antigen surface structure. Mutations are highlighted in green (K417N), cyan (E484K) and yellow (N501Y). (**B**) Computation modelling for PHH-1V vaccine. Top: AlphaFold2 results for the B.1.351-B.1.1.7 construct. This highlights the presence of two different construct conformations: (i) stabilized N-/C-terminal conformation, and (ii) adopting protein-protein interactions. Bottom: hACE2 receptor-construct model derived from MD calculations of the B.1.351-B.1.1.7 construct. RBD residues 1 to 219 and 220 to 439 are shown in grey and orange, respectively, whereas ACE2 monomers are shown as a transparent surface and cartoon representation in violet and purple. (**C**) SDS-PAGE. The reduced and non-reduced purified antigens were loaded at three serial dilutions: 1/10, 1/20 and 1/40. M: molecular weight ladder. C: BSA control. (**D**) SEC-HPLC chromatogram of the purified antigen. (**E**) Surface plasmon resonance (SPR) for the quantitative evaluation of the affinity between the antigen and its natural ligand, the human ACE2 receptor. RU: resonance units.

**Figure 2.**
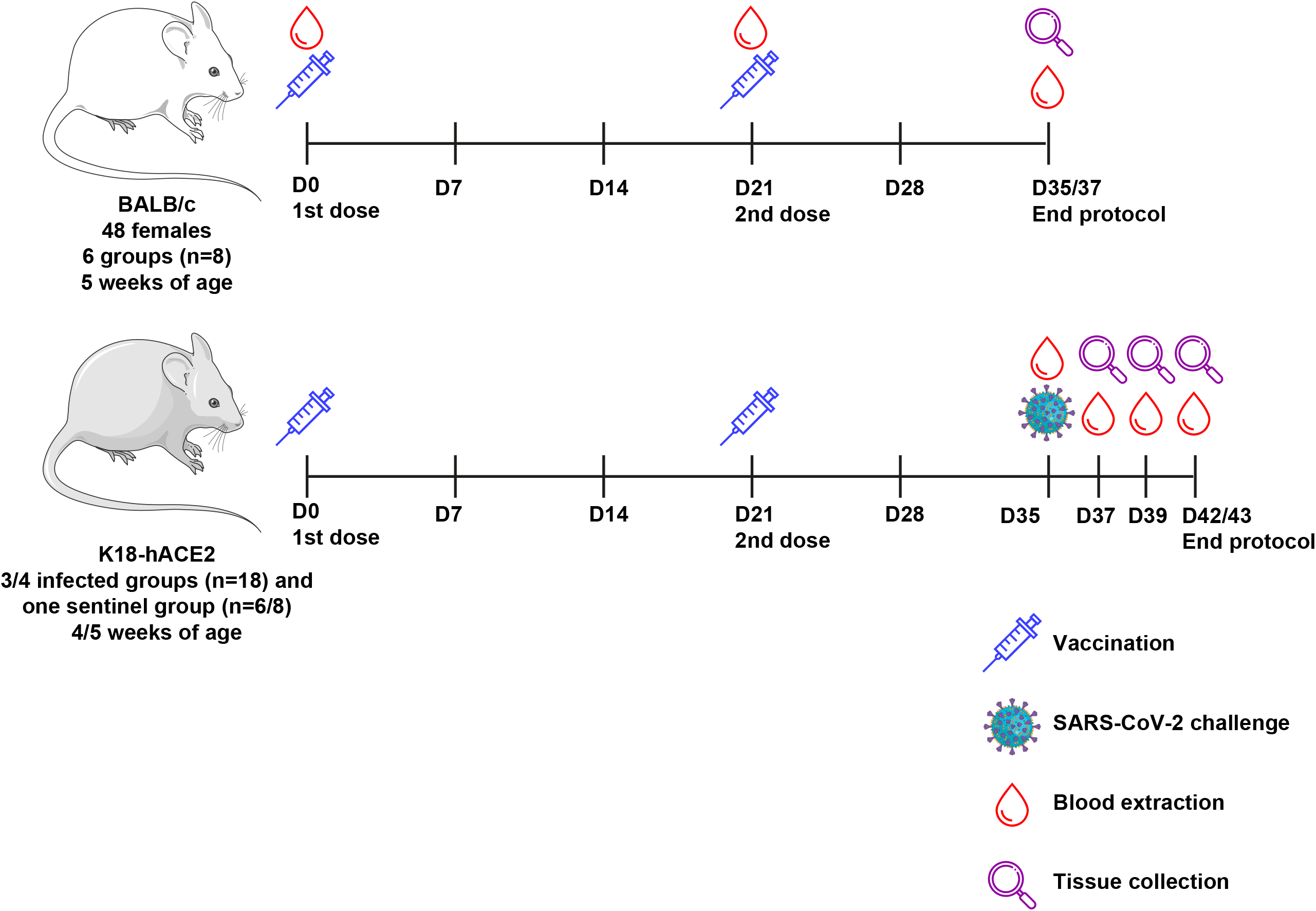
Schematic representation of the experimental protocol in BALB/c and K18-hACE2 mice for the safety, immunogenicity, and efficacy assessment. For safety and immunogenicity assays (in the top side), forty-eight 5-week-old female BALB/c mice were allocated to 6 groups (n=8) and were injected intramuscularly with two doses of 0.1 mL of the PHH1-1V vaccine on days 0 (prime) and 21 (boost). Then, animals were monitored daily for clinical signs and bodyweight was recorded weeKly until D35/D37, when they were euthanised and both spleens and blood were collected. For safety, immunogenicity and efficacy assays (in the bottom side), K18-hACE2 mice were allocated to 4 groups (efficacy against SARS-CoV-2 D614G) or 3 groups (efficacy against different VoCs), and were injected intramuscularly with two doses of 0.1 mL of the PHH1-1V vaccine on days 0 (prime) and 21 (boost). On D35 animals were challenged with 10^3^ TCID_50_ of the SARS-CoV-2 or different VoCs, blood samples were collected to analyse neutralising activity, and they were monitored daily for clinical signs and mortality. Then, challenged animals were chronologically euthanised on D37, D39 and D42/D43; and several tissue samples were collected for several analyses. Schematic artwork used in this figure is provided by Servier Medical Art under a Creative Commons Attribution 3.0 Unported License.

The protein-protein interaction energies of two construct variant candidates, B.1.351-B.1.1.7 and B.1.1.7-B.1.351, were estimated by means of molecular mechanics-generalised Born surface area (MM-GBSA) calculations. MM-GBSA results highlighted the B.1.351-B.1.1.7 variant as the preferred construct candidate with a protein-protein interaction energy of −78.23 kcal·mol^-1^ as compared to the value of -98.06 kcal·mol^-1^ for the inverse construct. This suggests that the N-/C-terminal stabilised conformation is energetically more favourable in the B.1.351-B.1.1.7 construct than in the B.1.1.7-B.1.351 construct. The interaction energies of both studied constructs, in a stabilised N-/C-terminal conformation, and the hACE2 receptor were also computed by means of MM-GBSA simulations (**Figure 2B**). Although both models showed similar binding affinities to hACE2, the Beta N-terminus plus Alpha C-terminus configuration clearly exposed those mutations involved in the higher affinity towards the human ACE2 receptor on the protein surface and the potential immune evasion by both variants. Hence, the selection of the B.1.351-B.1.1.7 fusion heterodimer as the PHH-1V vaccine antigen was based on the lower free energy required for the formation of the stabilised N-/C-terminal conformation.

The heterodimer is expressed in mammalian CHO cells and is formulated with the SQBA adjuvant. After expressing the antigen in a bioreactor fed-batch cultivation, it is purified by a downstream process consisting of sequential stages, including depth and tangential filtration, chromatography steps, and sterile filtrations. The final product is a highly purified antigen, as determined by SDS-PAGE (**Figure 1C**) and SEC-HPLC (**Figure 1D**), suitable for vaccine formulation. Moreover, surface plasmon resonance (SPR) analysis showed an affinity constant of 0.099 nM for hACE2 (**Figure 1E**).

### 2.2. Recombinant RBD fusion heterodimer antigen immunogenicity in BALB/c mice

#### 2.2.1. RBD-specific binding and neutralising antibody titres upon PHH-1V vaccination

BALB/c mice (Environ, IN, USA) were immunised with different doses of the recombinant RBD fusion heterodimer antigen (group B: 0.04 μg, group C: 0.2 μg, group D: 1 μg, group E: 5 μg; and group F: 20 μg) on days (D) 0 and 21. Mice were also immunised with PBS as a control group (group A). A schematic view of the immunisation protocol is depicted in **Figure 2**.

The prime immunisation of BALB/c mice with the PHH-1V candidate induced higher titres of RBD binding antibodies in groups C to F compared to the control (group A, immunised with PBS) on day 21 post-first immunisation (D21) (*p<0.01*) (**Figure 3A**). After the prime-boost immunisation, all vaccinated groups (B to F) reached higher IgG titres than the control group on D35/D37 (14/16 days after the boost; *p<0.01*). On D35/D37, specific SARS-CoV-2 RBD-binding antibodies were detected in groups B to D in a dose-dependent manner, with significant differences between these groups (*p<0.01*). However, no significant differences were observed between the groups immunised with more than 1 μg of recombinant RBD fusion heterodimer antigen (groups D to F). Therefore, the IgG response was saturated from 1- μg immunisation. Likewise, the IgG2a/IgG1 ratios were calculated as a surrogate of the Th1/Th2 cellular immune response to estimate the type of cellular immune response elicited by the vaccine. The IgG2a/IgG1 ratio of groups E and F was 0.74 and 0.75, respectively, which suggests a balanced Th1/Th2 immunogenic response upon PHH-1V vaccination in mice (**Figure 3B**).

**Figure 3.**
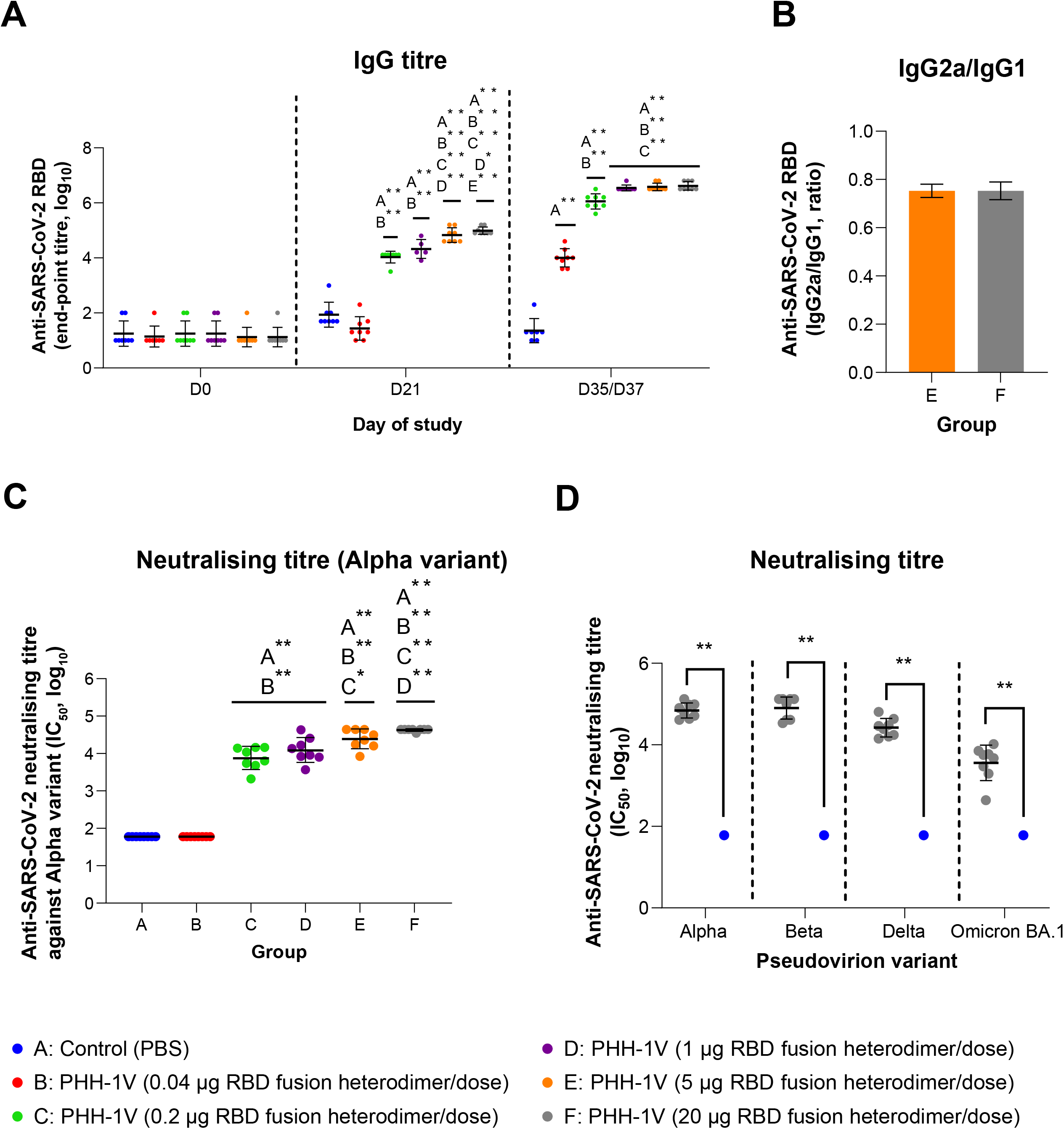
Analysis of the antibody response upon PHH-1V vaccination in mice. (**A**) SARS-CoV-2 RBD-specific IgG responses in groups A to F on days D0, D21 and D35/D37. Endpoint antibody titres determined by ELISA in female BALB/c mice are shown. Log_10_ IgG titres were analysed by means of a linear mixed effects model. (B) Endpoint titre ratios of IgG2a to IgG1 in female mice vaccinated with PHH-1V vaccine (groups E and F). Analyses of IgG1 and IgG2 subclasses in groups E and F were performed by ELISA on serum samples taken on day D35/D37. Data was analysed by means of a Mann-Whitney U-test. (C) Neutralising antibody responses in groups A to F. SARS-CoV-2 neutralising antibody titres in sera, against pseudoviruses that express the S protein with the Alpha sequence, were determined by PBNA 14/16 days after the second dose of each vaccine (D35/D37). Sera from female BALB/c mice collected on D35/D37 were assessed for pseudovirus-neutralising activity.Log_10_ IC_50_ data was analysed using a generalized least squares (GLS) model, employing one-sample tests against the null H_0_: μ = 1.78 for comparison of estimated marginal mean against groups A and B. (D) Neutralising antibody responses against multiple SARS-CoV-2 variants (Alpha, Beta, Delta, Omicron BA.1) by PBNA upon 20-μg RBD fusion heterodimer/dose immunization. Sera mice from groups A and F collected on D35/D37 were assessed for pseudovirus-neutralising activity as pool sera or individual sera, respectively. For the analysis of this data, one-sample t-tests against the null H_0_: μ = 1.78 were employed. Each data point represents an individual mouse serum, with bars representing the mean titre per group ± SD. Statistically significant differences between groups are indicated with a line on top of each group: * *p<0.05*; ** *p<0.01*; ^+^ *0.05<p<0.1*.

SARS-CoV-2 neutralising antibodies titres in sera from BALB/c mice were determined by a pseudovirus-based neutralisation assay (PBNA) against the S protein of different variants on D35/D37 (14/16 days after the boost). Prime-boost immunisation of groups C to F induced higher neutralising antibody titres against the S protein of the Alpha variant compared to the control group A (*p<0.01*) (**Figure 3C**). No neutralising antibody response was observed in group B, although IgG binding antibodies were detected on D35/D37. The mean neutralising antibody titres observed in groups C and D remained the same since no statistically significant differences were observed. However, vaccination with 5 μg (group E) and 20 μg (group F) of RBD fusion heterodimer antigen induced higher neutralising titres than group C and groups C and D, respectively. Interestingly, high neutralising titres against all the tested variants (Alpha, Beta, Delta and Omicron BA.1) were detected in sera from group F compared to control group A (*p<0.01*) (**Figure 3D**).

#### 2.2.2. RBD-specific cellular immune response upon PHH-1V vaccination

The characterisation of the antigen-specific response of splenic T cells 14/16 days after the boost immunisation was performed by intracellular cytokine staining (ICS) and enzyme-linked immunospot (ELISpot) assays in female BALB/c mice from groups A (control), E and F (vaccinated with 5 μg or 20 μg of recombinant protein RBD fusion heterodimer, respectively). The ICS data indicate that upon stimulation with an RBD peptide pool, splenocytes from group F displayed significant activation of CD4^+^ T cells expressing IFN-γ (*p<0.01* IL-2 (*p<0.05*) and Th1-like cytokines (IFN-γ and/or TNF-α and/or IL-2; *p<0.05*) compared to the control group (**Figure 4A**). No significant antigen-specific response of CD4^+^ T cells expressing TNF-α or IL-4 was observed in group F when compared to the control group. Notably, immunisation of mice with a lower RBD dose (group E) did not induce CD4^+^ T cells secreting IFN-γ, TNF-α, IL-2 and/or IL-4 after the *in vitro* restimulation. Furthermore, splenocytes from group F showed activation of CD8^+^ T cells expressing IFN-γ (*0.05<p<0.1*) and IL-2 (*p<0.05*) after the antigen-specific restimulation compared to the control group (**Figure 4B**). Likewise, splenocytes from group F also elicited significantly higher CD8^+^ T cells expressing IFN-γ (*p<0.05*) and IL-2 (*p<0.01*) compared to the group E. No CD8^+^ T cell response was observed in splenocytes from group E compared to the control group.

**Figure 4.**
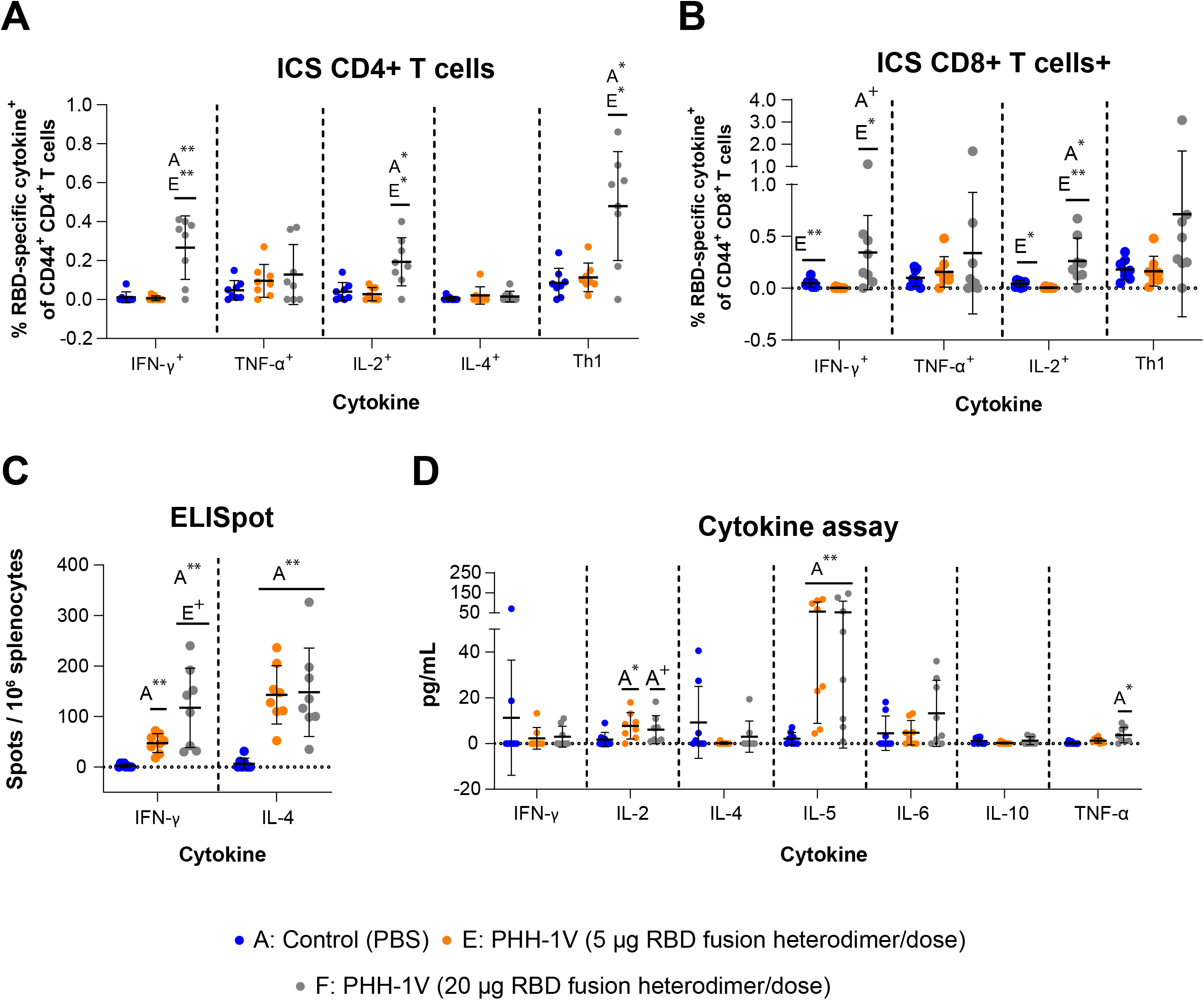
PHH-1V-induced CD4^+^ and CD8^+^ T cell responses, and extracellular cytokine levels in vaccine-induced splenocytes from mice. Splenocytes from vaccinated female BALB/c mice were isolated 14/16 days after boost immunization (D35/D37), stimulated with RBD peptide pools, and analysed by intracellular cytokine staining. The frequencies of cytokine expressing CD4^+^ T cells (A) and CD8^+^ T cells (B) are shown. The frequencies of CD4^+^ and CD8^+^ T cells expressing Th1 cytokines (sum of IFN-γ, TNF-α, IL-2) are also shown. The cytokine expression in splenocytes stimulated with the medium was considered the background value and this was subtracted from peptide-specific responses. Data were analysed using a generalized least squares (GLS) model on the arcsine-square root-transformed percentage values. (**C**) Splenocytes from vaccinated BALB/c mice were isolated 14/16 days after boost immunization (D35/D37), stimulated with RBD peptide pools, and analysed by IFN-γ and IL-4-specific ELISpot assays. Data were analysed using a generalized least squares (GLS) model on the arcsine-square root-transformed percentage values. (D) Extracellular cytokines were measured by Luminex Multiplex in supernatants from BALB/c splenocytes stimulated with a pool of peptides from SARS-CoV-2 RBD. Cytokine levels in splenocytes stimulated with the medium were considered the background value and these were subtracted from the responses measured from the RBD peptide pool for each individual mouse. Data were analysed using Kruskal-Wallis’ H test and Dunn’s post-hoc with Holm’s correction for multiple testing or Mann-Whitney’s U-test. Each data point represents an individual mouse value, with bars representing the mean ± SD. Statistically significant differences between groups are indicated with a line on top of each group: * *p<0.05*; ** *p<0.01*; ^+^ *0.05<p<0.1*.

The IFN-γ and IL-4 ELISpot assays showed no significant differences between the two doses of recombinant protein RBD fusion heterodimer (group E vs group F) (**Figure 4C**).However, both groups showed a higher percentage of IFN-γ^+^ and IL-4^+^ spots compared to the control group (*p<0.01*). Importantly, the percentage of IFN-γ^+^ and IL-4^+^ in group F was similar, denoting a balanced Th1/Th2 response, while the percentage of IL-4^+^ spots was significantly higher than IFN-γ^+^spots in group E (*p<0.01*), suggesting a Th2-biased response in mice immunised with 5 μg of recombinant protein RBD fusion heterodimer.

Extracellular cytokine levels were measured by Luminex Multiplex in supernatants from splenocytes stimulated with a pool of peptides from SARS-CoV-2 RBD. The levels of IL-2 (*p<0.05*) and IL-5 (*p<0.01*) were higher in the supernatants from group E splenocytes compared to the control group (**Figure 4D**). Similarly, the levels of IL-5 (*p<0.01*) and TNF-α (p<0.05) were statistically higher in group F compared to group A. A tendency towards an increase in the levels of IL-2 (*0.05<p<0.1*) was also observed in group F compared to group A.

### 2.3. Recombinant RBD fusion heterodimer antigen immunogenicity and efficacy against SARS-CoV-2 Wuhan/D614G in K18-hACE2 mice

To analyse the immunogenicity and protective efficacy of the PHH-1V vaccine candidate against COVID-19 and the pathogenic outcomes derived from the SARS-CoV-2 infection, the mouse strain B6.Cg-Tg(K18-ACE2)2Prlmn/J (K18-hACE2) (Jackson Laboratories, ME, USA) was used as a challenge model. Groups were vaccinated intramuscularly with PBS (groups A and B), 10 μg of PHH-1V (group C) or 20 μg of PHH-1V (group D) following the two-dose prime-and- boost schedule: 1^st^ dose (prime) on D0 and 2^nd^ dose (boost) on D21 (**Figure 2**).The SARS-CoV-2 (Wuhan/D614G strain) challenge was performed on group B-D animals on D35 through intranasal instillation.

The primary endpoint for reporting the protective capacity of the vaccine candidates was weight loss and/or mortality post-challenge. Clinical signs and survival are presented in **Figure 5A**.Clinical signs of the SARS-CoV-2 infection were observed only in the non-vaccinated and infected group (B) on days 5 (3 animals) and 6 (3 animals) post-challenge. In all cases, clinical signs led to endpoint criteria and the animals were euthanised. Thus, survival of group B was significantly different than the other groups (*p<0.01*). The daily individual bodyweights of each group during the vaccination period and post-challenge are shown in **Figures S1 B** and **5B**, respectively. The animals from group B experienced remarkable weight loss from D3 post-challenge onwards, as expected due to the SARS-CoV-2 infection, showing a significantly lower weight compared to vaccinated animals from groups C and D on D5 and 6 post-challenge (*p<0.01*).

**Figure 5.**
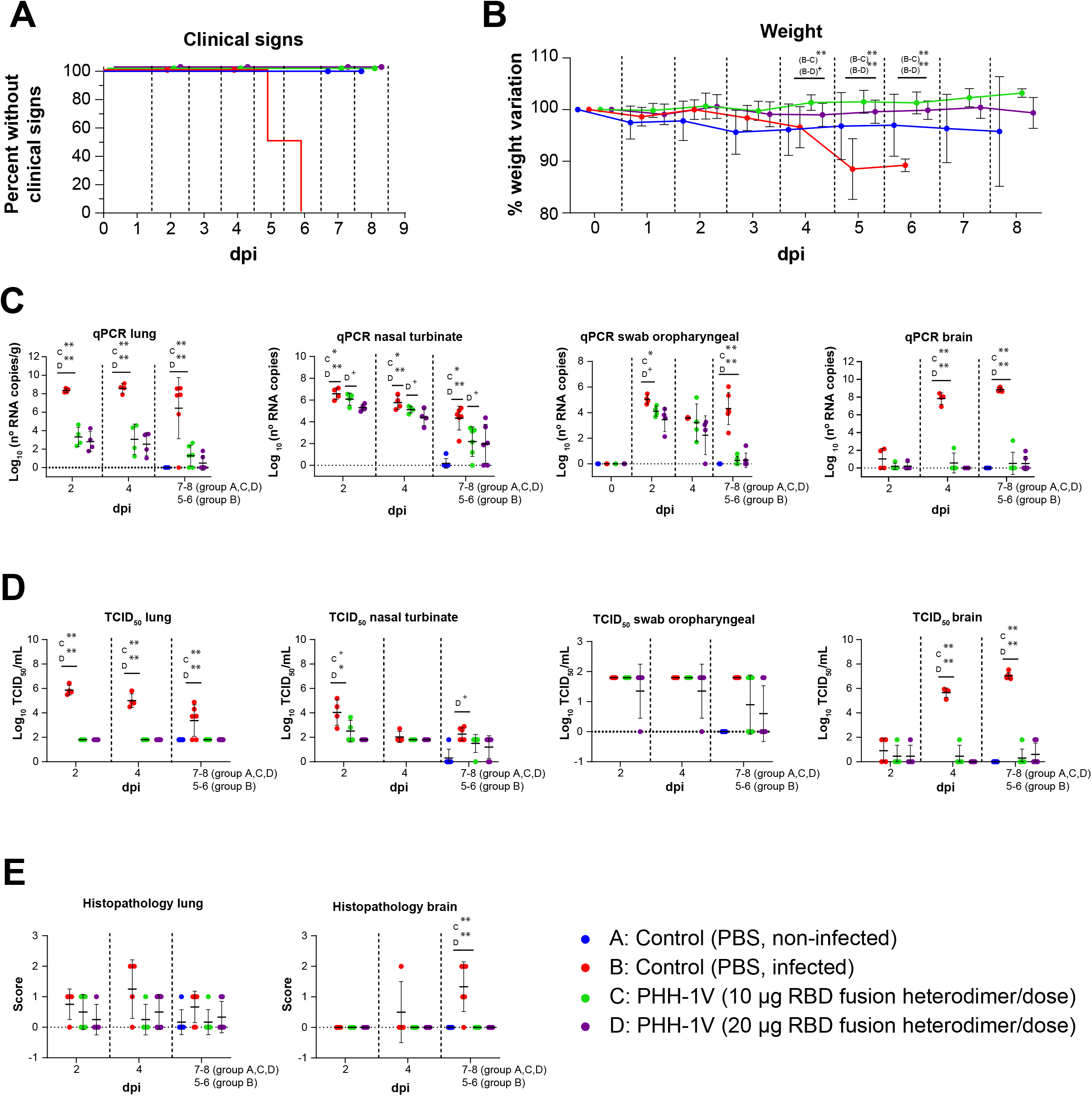
Protective efficacy of PHH-1V vaccine in K18-hACE2 mice upon SARS-CoV-2 challenge. Group A (n=8, 4F + 4M), group B (n=l8, 9F + 9M), group C (n=18, 9F + 9M), and group D (n=18, 9F + 9M). (**A**) Survival curves for groups of immunized K18-hACE2 mice with PHH-1V vaccine and control groups. Survival analysis (Kaplan-Meier estimates and log-rank test to compare groups) was performed to study differences in time to/before clinical signs and mortality. (B) Mean weight change after SARS-CoV-2 challenge calculated as a percentage of the pre-challenge weight in K18-hACE2 mice. A linear mixed effects model on the body weight change data was performed considering groups B, C and D. Points represent the average weight variation in each group and error bars depict a +/-SD interval. (**C**) SARS-CoV-2 RT-qPCR (number of copies) in the lungs, nasal turbinate, oropharyngeal swabs and brain collected from challenged animals. (D) Viral titres were determined using a standard TCID_50_ assay on positive samples of RT-qPCR (in some exceptional cases, RT-qPCR and viral isolation were performed in parallel for logistical reasons). RT-qPCR-negative samples are represented as 0 TCID_50_/mL. The detection limit was set at 1.8 TCID_50_/mL. (E) Histopathological analyses from the lungs and brain were determined for all animals. For each tissue sample, lesions were classified as follows: multifocal broncho-interstitial pneumonia; multifocal lymphoplasmacytic rhinitis; multifocal lymphoplasmacytic meningoencephalitis; and multifocal mononuclear inflammatory infiltrates within and around muscular fibres. Lesions were evaluated with the following score: 0 (no lesion); 1 (mild lesion); 2 (moderate lesion); and 3 (severe lesion). Samples of groups A, C and D correspond to 2 (D37), 4 (D39) and 7 dpi (D42 for males) or 8 dpi (D43 for females); samples of group B were taken 2 (D37), 4 (D39), and 5 dpi (D40; n=3) or 6 dpi (D41; n=3), when animals reached the endpoint criteria. Generalized least squares models or Kruskal-Wallis and Dunn’s *post-hoc* tests were employed for the analysis of the RT-qPCR, TCID_50_ and histopathological data depending on verification of assumptions. Each data point represents an individual mouse value, with bars representing the mean ± SD. Statistically significant differences between groups are indicated with a line on top of each group: * *p<0.05; ** p<0.01; ^+^ 0.05<p<0.1*. DPI: days post infection. See also **Figures S1-S4**.

SARS-CoV-2 neutralising antibodies against the original Wuhan/D614G strain were also analysed in SARS-CoV-2 infected K18-hACE2 mice upon vaccination to study the immunogenicity of PHH-1V vaccine in a humanised mice model (**Figure S2 A**).Animals from group D elicited significant higher SARS-CoV-2-specifc neutralising titres 0-, 2- and 4-days post-infection (dpi) (*p<0.01*) and 7-8 dpi (*p<0.05*) compared to group B. Animals from group C elicited higher specific neutralising titres 0, 2 (*p<0.01*), 4 dpi (*p<0.05*) and 7-8 dpi (*p<0.01) vs*. control group (B). Furthermore, the levels of neutralising antibodies were similar between both vaccinated groups.

Total viral RNA was determined in the lungs, nasal turbinate, oropharyngeal swabs and brain (**Figure 5C**),but also in trachea, heart, pharynx and spleen (**Figure S3**).Viral RNA was determined by real-time quantitative polymerase chain reaction (RT-qPCR) on D37 (2 dpi), D39 (4 dpi), D42 (in males, 7 dpi) and D43 (in females, 8 dpi), or at the time of euthanasia in animals reaching endpoint criteria before the scheduled euthanasia day. Immunisation with 10 μg of PHH-1V (group C) reduced the viral load measured by PCR in the lungs on all dpi studied (*p<0.01*), in nasal turbinate on all dpi studied (*p<0.05*), in oropharyngeal swabs 2 dpi (*p<0.05*) and 7-8 dpi (*p<0.01*), and in brain 4 and 7-8 dpi (*p<0.01*) compared to the infected control (group B). Vaccination with 20 μg of PHH-1V (group D) also reduced the viral load measured by PCR in the lungs on all dpi studied (*p<0.01*), in nasal turbinate on all dpi studied (*p<0.01*), in oropharyngeal swabs 7-8 dpi (*p<0.01*), and in brain 4 and 7-8 dpi (*p<0.01*) compared to group B (**Figure 5C**). Likewise, both vaccinated groups reduced significantly viral RNA in the trachea (*p<0.01*), pharynx (*p<0.05*) and spleen (*p<0.01*) compared to control group (**Figure S3**). In heart, there was a tendency towards a decrease in both vaccinated groups 4 dpi compared to group B.

Virus titres were determined using a standard 50% tissue culture infectious dose (TCID_50_) assay on positive samples of RT-qPCR in lungs, nasal turbinates, oropharyngeal swabs and brain (**Figure 5D**).Samples from groups C and D had a significant lower infective viral load in the lungs during the entire post-challenge period (*p<0.01*) and in the brain 4 and 7-8 dpi (*p<0.01*). In the nasal turbinate, a significant lower infective viral load was observed 2 dpi in group D (*p<0.05*) compared with group B, and there was a tendency towards a decrease in group C (*0.05<p<0.1*) 2 dpi and group D (*0.05<p<0.1*) 7-8 dpi compared to group B.

Histopathological analyses were performed on lungs and brain of all animals (**Figure 5E**).Infected non-vaccinated animals (group B) had a higher histopathological score in the brain 7-8 dpi compared to group C and D (*p<0.01*). No significant differences between groups were observed in the histopathological score of the lungs, but these were numerically higher 4 dpi in group B compared to both vaccinated groups. To support the histopathological scores of **Figure 5E**, we also chose representative sections of brain and lung from study mice showing scores of 0 (lack of lesions), 1 (mild lesions), and 2 (moderate lesions) (**Figure S4**).None of the animals of the study showed lesions with score 3 (severe lesions). Furthermore, other tissues such as spleen, trachea or heart were analysed and no lesions were found in any of the studied animals (**Figure S5 A-C**).

### 2.4. Recombinant RBD fusion heterodimer antigen immunogenicity and efficacy against different SARS-CoV-2 VoCs in K18-hACE2 mice

We next assessed the protective efficacy of PHH-1V vaccine against Beta, Delta and Omicron BA.1 SARS-CoV-2 VoCs in the K18-hACE2 mice model. Animals were vaccinated intramuscularly with two doses of PBS (groups A and B) or 20 μg of PHH-1V vaccine (group C) on D0 and D21, and then animals from groups B and C were intranasally challenged on D35 with Beta, Delta or Omicron BA.1 SARS-CoV-2 variants. Animals were monitored for weight loss and mortality for 7 days and then were euthanised at 2, 4 and 7 dpi to analyse viral load in oropharyngeal swabs and lungs by both RT-qPCR and viral titration, and also histopathology in lung sections (**Figure 6–8**).

**Figure 6.**
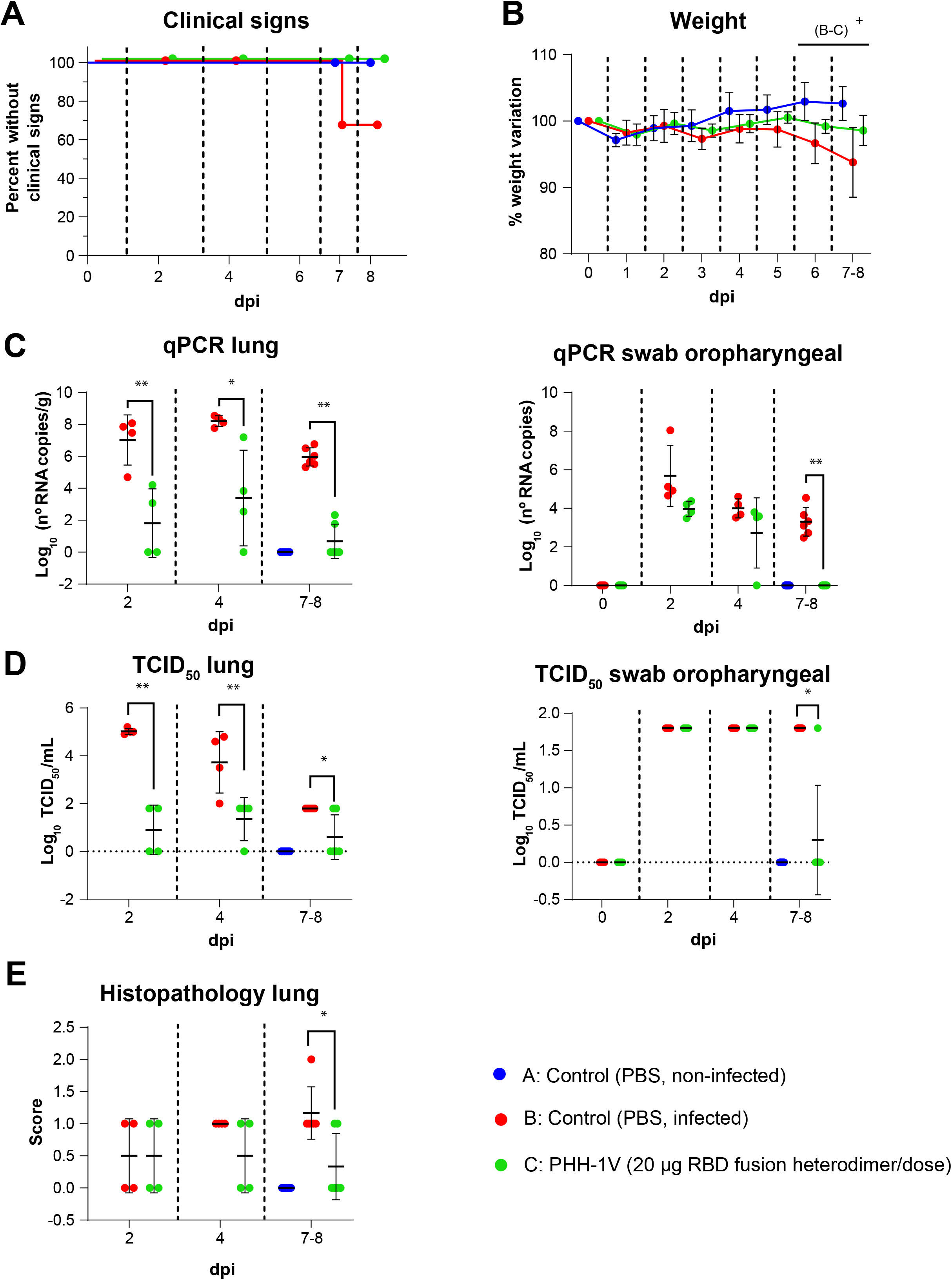
Protective efficacy of PHH-1V vaccine in K18-hACE2 mice upon challenge with SARS-CoV-2 Beta variant. Group A (n=8, 4F + 4M), group B (n=18, 9F + 9M), and group C (n=18, 9F + 9M). (**A**) Survival curves of animals from PHH-1V vaccinated groups and non-vaccinated groups. Survival analysis (Kaplan-Meier estimates and log-rank test to compare groups) was performed to study differences in time to/before clinical signs and mortality. (**B**) Mean weight change after Beta variant challenge calculated as a percentage of the pre-challenge weight in K18-hACE2 mice. A linear mixed effects model on the body weight change data was performed considering groups B and C. Points represent the average weight variation in each group and error bars depict a +/- SD interval. (**C**) SARS-CoV-2 RT-qPCR (number of copies) in the lungs and oropharyngeal swabs collected from challenged animals. (**D**) Viral titres were determined using a standard TCID_50_ assay on positive samples of RT-qPCR. Negative samples are represented as 0 TCID_50_/mL. The detection limit was set at 1.8 TCID_50_/mL. (**E**) Histopathological analyses from the lungs were determined for all animals. For each tissue sample, lesions were classified as follows: multifocal broncho-interstitial pneumonia; multifocal lymphoplasmacytic rhinitis; multifocal lymphoplasmacytic meningoencephalitis; and multifocal mononuclear inflammatory infiltrates within and around muscular fibres. Lesions were evaluated with the following score: 0 (no lesion); 1 (mild lesion); 2 (moderate lesion); and 3 (severe lesion). All the samples correspond to 2 (D37), 4 (D39) and 7 dpi (D42 for males) or 8 days post infection (D43 for females); or at the time of euthanasia in animals reaching endpoint criteria before the scheduled euthanasia day. Generalized least squares models or Mann-Whitney tests were employed for the analysis of the RT-qPCR, TCID_50_ and histopathological data depending on verification of assumptions. Each data point represents an individual mouse value, with bars representing the mean ± SD. Statistically significant differences between groups are indicated with a line on top of each group: *\* p<0.05; ** p<0.01; ^+^ 0.05<p<0.1*. DPI: days post infection. See also **Figure S2**.

**Figure 7.**
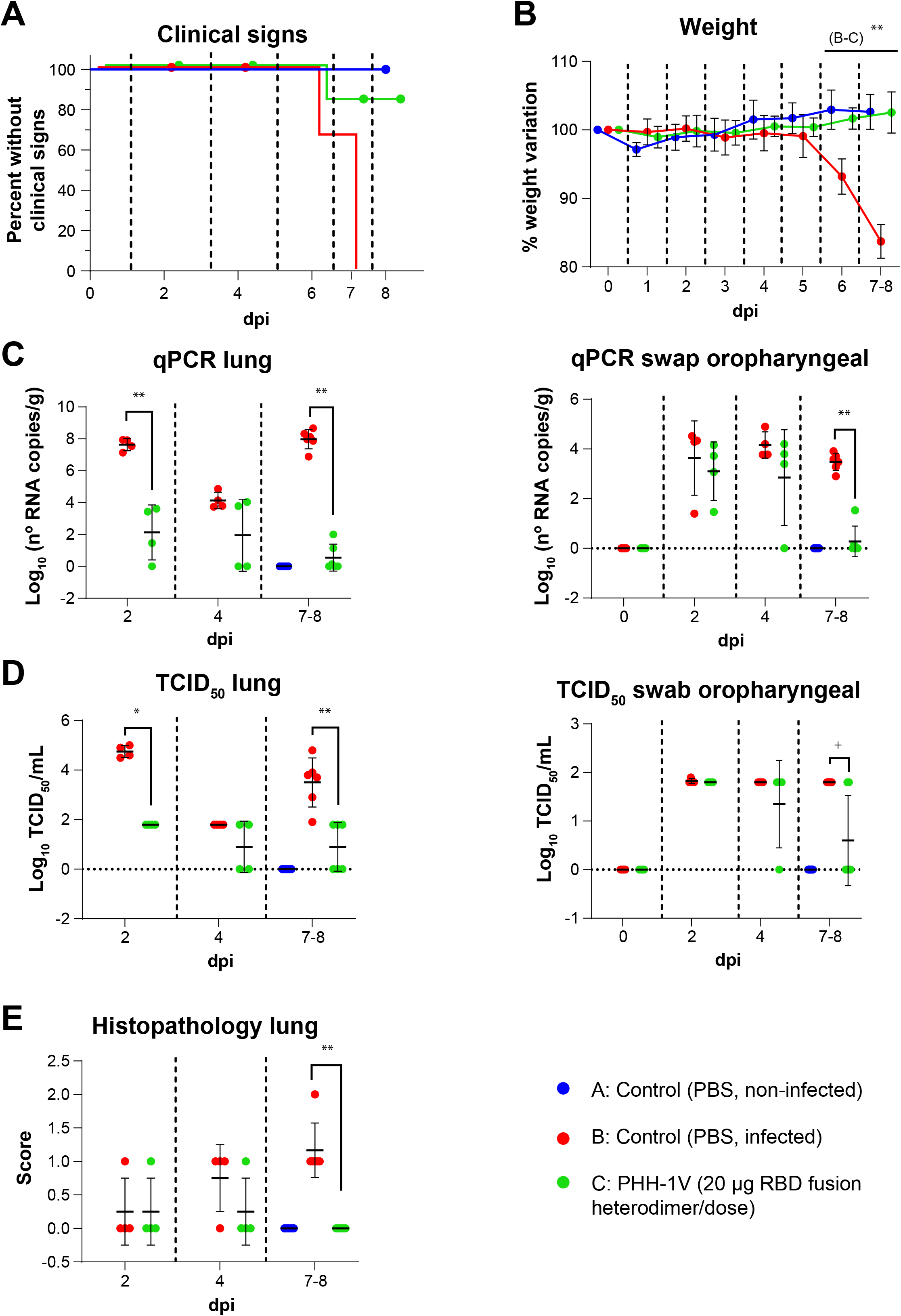
Protective efficacy of PHH-1V vaccine in K18-hACE2 mice upon challenge with SARS-CoV-2 Delta variant. Group A (n=8, 4F + 4M), group B (n=18, 9F + 9M), and group C (n=18, 9F + 9M), (**A**) Survival curves of animals from PHH-1V vaccinated groups and non-vaccinated groups. Survival analysis (Kaplan-Meier estimates and log-rank test to compare groups) was performed to study differences in time to/before clinical signs and mortality. (**B**) Mean weight change after Delta variant challenge calculated as a percentage of the pre-challenge weight in K18-hACE2 mice. A linear mixed effects model on the body weight change data was performed considering groups B and C. Points represent the average weight variation in each group and error bars depict a +/- SD interval. (**C**) SARS-CoV-2 RT-qPCR (number of copies) in the lungs and oropharyngeal swabs collected from challenged animals. (**D**) Viral titres were determined using a standard TCID_50_ assay on positive samples of RT-qPCR. Negative samples are represented as 0 TCID_50_/mL. The detection limit was set at 1.8 TCID_50_/mL. (**E**) Histopathological analyses from the lungs were determined for all animals. For each tissue sample, lesions were classified as previously assays. Lesions were evaluated with the following score: 0 (no lesion); 1 (mild lesion); 2 (moderate lesion); and 3 (severe lesion). All the samples correspond to 2 (D37), 4 (D39) and 7 dpi (D42 for males) or 8 days post infection (D43 for females); or at the time of euthanasia in animals reaching endpoint criteria before the scheduled euthanasia day. Generalized least squares models or Mann-Whitney tests were employed for the analysis of the RT-qPCR, TCID_50_ and histopathological data depending on verification of assumptions. Each data point represents an individual mouse value, with bars representing the mean ± SD. Statistically significant differences between groups are indicated with a line on top of each group: * *p<0.05; ** p<0.01; ^+^ 0.05<p<0.1*. DPI: days post infection. See also **Figure S2**.

**Figure 8.**
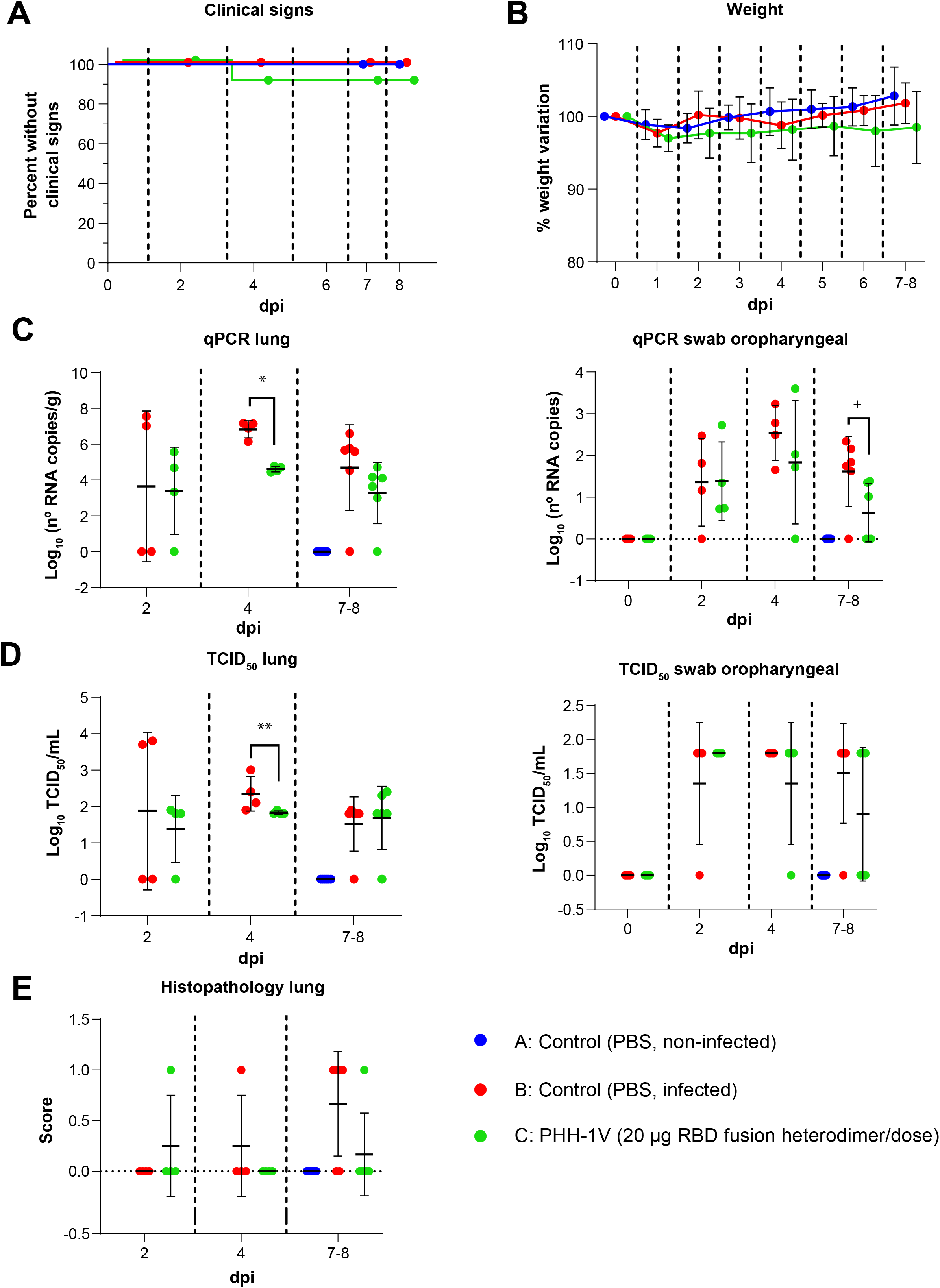
Protective efficacy of PHH-1V vaccine in K18-hACE2 mice upon challenge with SARS-CoV-2 Omicron BA.1 variant. Group A (n=8, 4F + 4M), group B (n=18, 9F + 9M), and group C (n=18, 9F + 9M), (**A**) Survival curves of animals from PHH-1V vaccinated groups and non-vaccinated groups. Survival analysis (Kaplan-Meier estimates and log-rank test to compare groups) was performed to study differences in time to/before clinical signs and mortality. (**B**) Mean weight change after Omicron BA.1 variant challenge calculated as a percentage of the pre-challenge weight in K18-hACE2 mice. A linear mixed effects model on the body weight change data was performed considering groups B and C. Points represent the average weight variation in each group and error bars depict a +/- SD interval. (**C**) SARS-CoV-2 RT-qPCR (number of copies) in the lungs and oropharyngeal swabs collected from challenged animals. (**D**) Viral titres were determined using a standard TCID_50_ assay on positive samples of RT-qPCR. Negative samples are represented as 0 TCID_50_/mL. The detection limit was set at 1.8 TCID_50_/mL. (**E**) Histopathological analyses from the lungs were determined for all animals. For each tissue sample, lesions were classified as previously assays. Lesions were evaluated with the following score: 0 (no lesion); 1 (mild lesion); 2 (moderate lesion); and 3 (severe lesion). All the samples correspond to 2 (D37), 4 (D39) and 7 dpi (D42 for males) or 8 days post infection (D43 for females); or at the time of euthanasia in animals reaching endpoint criteria before the scheduled euthanasia day. Generalized least squares models or Mann-Whitney tests were employed for the analysis of the RT-qPCR, TCID_50_ and histopathological data depending on verification of assumptions. Each data point represents an individual mouse value, with bars representing the mean ± SD. Statistically significant differences between groups are indicated with a line on top of each group: * *p<0.05*; ** *p<0.01*; ^+^ *0.05<p<0.1*. DPI: days post infection. See also **Figure S2**.

#### 2.4.1. Immunogenicity and efficacy of PHH-1V against SARS-CoV-2 Beta variant (B.1.351)

Clinical signs of the SARS-CoV-2 infection were observed only in two animals from the non-vaccinated and infected group (group B) on day 7 post-challenge. Clinical signs curves are shown in **Figure 6A**. Animals from group B had a lower weight (*0.05<p<0.1*) 6 and 7-8 dpi than animals vaccinated with PHH-1V (group C) (**Figure 6B**).

SARS-CoV-2 neutralising antibodies against Beta variant were also analysed upon vaccination with PHH-1V in infected K18-hACE2 mice. PHH-1V vaccination elicited higher neutralising titres against Beta variant 0 dpi (D35, pre-challenge) (*p<0.05*) and 7-8 dpi (*p<0.01*) compared to infected control animals (**Figure S2 B**).

Viral load was determined in lungs and oropharyngeal swabs by RT-qPCR and TCID_50_ on D37 (2 dpi), D39 (4 dpi) and D42 (in males, 7 dpi) or D43 (in females, 8 dpi), or at the time of euthanasia in animals reaching endpoint criteria before the scheduled euthanasia day. Immunisation with 20 μg of PHH-1V (group C) reduced the viral RNA in the lungs 2 dpi (p<0.01), 4 dpi (p<0.05) and 7-8 dpi (*p<0.01*), but also in oropharyngeal swabs 7-8 dpi (p<0.01) compared to the infected control (group B) (**Figure 6C**). Similarly, PHH-1V vaccination reduced the infectious viral load in lungs 2 dpi (*p<0.01*), 4 dpi (*p<0.01*) and 7-8 dpi (*p<0.05*), and also in oropharyngeal swabs 7-8 dpi (*p<0.05*) compared to the infected control (**Figure 6D**).

Histopathological scores were also calculated in lung sections from all study animals. Infected control animals had a higher histopathological score in the lung 7-8 dpi compared to PHH-1V vaccinated animals (*p<0.05*).

#### 2.4.2. Immunogenicity and efficacy of PHH-1V against SARS-CoV-1 Delta variant (B.1.617.2)

Clinical signs were observed in all non-vaccinated and infected animals (group B) 6 dpi (2 animals) or 7 dpi (4 animals). Three of the six animals showing clinical signs in group B reached the endpoint criteria and were euthanised. In contrast, only one animal of group C showed clinical signs 6 dpi (appearance alteration of score 1). Therefore, curves were significantly different between both groups (B vs C; *p<0.05*) (**Figure 7A**). Weight variation was also significantly lower in group B compared to group C from 6 dpi onwards (*p<0.01*) (**Figure 7B**).

Neutralising titres induced by PHH-1V vaccine were measured against Delta variant 0 dpi (pre-challenge) and 7-8 dpi (post-challenge). PHH-1V vaccination elicited higher neutralising titres against Delta variant 0 dpi (D35, pre-challenge) and 7-8 dpi (*p<0.01*) compared to infected control animals (group B) (**Figure S2 C**).

Viral load was also determined in lungs and oropharyngeal swabs by RT-qPCR and TCID_50_ on D37 (2 dpi), D39 (4 dpi) and D42 (in males, 7 dpi) or D43 (in females, 8 dpi), or at the time of euthanasia in animals reaching endpoint criteria before the scheduled euthanasia day. Immunisation with 20 μg of PHH-1V (group C) reduced the viral RNA in the lungs 2 dpi and 7-8 dpi (*p<0.01*), and in oropharyngeal swabs 7-8 dpi (p<0.05) compared to the infected control group (**Figure 7C**).Likewise, PHH-1V vaccination reduced the infectious viral load in lungs 2 dpi (p<0.05) and 7-8 dpi (*p<0.01*) compared to the infected control group (**Figure 7D**). There was a decreasing trend in the infectious viral load from oropharyngeal swabs of group C 7-8 dpi compared to group B (*0.05<p<0.1*). Furthermore, infected control animals had a higher histopathological score in the lung 7-8 dpi compared to PHH-1V vaccinated animals (*p<0.01*) (**Figure 7E**).

#### 2.4.3. Immunogenicity and efficacy of PHH-1V against SARS-CoV-1 Omicron variant (BA.1)

Additionally, we tested the immunogenicity and efficacy of PHH-1V vaccine against Omicron BA.1, the predominant variant at the time these assays were conducted. No significant differences were observed between groups in clinical signs curves (**Figure 8A**) and weight loss (**Figure 8B**). However, PHH-1V vaccinated animals (group C) showed significantly less viral RNA in lungs 4 dpi (*p<0.05*) (**Figure 8C**) and less infectious SARS-CoV-2 in lungs 4 dpi (*p<0.01*) (**Figure 8D**). There was also a decreasing trend in the viral RNA from oropharyngeal swabs of group C 7-8 dpi compared to group B (*0.05<p<0.1*).

Furthermore, neutralising titres induced by PHH-1V vaccine were measured against Omicron BA.1 variant 0 dpi (D35, pre-challenge) and 7-8 dpi (D42-D43, during post-challenge). PHH-1V vaccination elicited higher neutralising titres against Omicron variant 0 dpi and 7-8 dpi (*p<0.01*) compared to infected control animals (group B) (**Figure S2 D**).

### 2.5. Safety of the recombinant RBD fusion heterodimer antigen after vaccination

The preclinical safety of the PHH-1V candidate vaccine was evaluated in BALB/c mice immunised with different doses of the RBD fusion heterodimer by measuring the bodyweight of each animal once a week until D35/D37. For additional safety information, clinical signs and local reactions were monitored. Differences in bodyweight were observed between the control group and some of the vaccinated groups at different times. The fact that the highest dose (group F) did not show significant differences with the control group during the entire study suggests that these differences in bodyweight are not related to the antigen composition. On the other hand, the differences observed in bodyweight cannot be attributed to the adjuvant because all the PHH-1V vaccines contain the same amount of adjuvant. The mean bodyweight of the control group was above the rest of the groups from the beginning of the study, which could have contributed to the differences observed with the rest of the groups throughout the study (**Figure S1 A**). No clinical signs or local reactions were detected after the vaccinations in BALB/c mice.

Additionally, the safety of PHH-1V was evaluated in humanised K18-hACE2 mice, and bodyweight and clinical signs were also monitored during the vaccination period. No significant changes in bodyweight were observed between the different groups (**Figure S1 B**), and vaccinated animals did not show clinical signs or local reactions. The histological evaluation of the injection site revealed a mild lesion (with multifocal mononuclear inflammatory infiltrates within and around muscular fibres) in one of the hind limbs of 1 animal vaccinated with the 10-μg RBD fusion heterodimer/dose on D2 and 2 animals vaccinated with the 20-μg RBD fusion heterodimer/dose on D4 post-challenge (**Figure S5 D**).

## 3. Discussion

In this study, the effect of a recombinant protein RBD fusion heterodimer dose on the immunogenicity and safety of the PHH-1V vaccine was tested in BALB/c and K18-hACE2 mice. Furthermore, the preclinical efficacy of the vaccine candidate was also assessed in SARS-CoV-2-infected and non-infected K18-hACE2 mice, as well as cross-protection induced against Beta, Delta and Omicron BA.1 VoCs. We show that the active substance of the PHH-1V vaccine candidate, the RBD fusion heterodimer, is stable and has an affinity constant of 0.099 nM against the human ACE2 receptor, which indicates an outstanding binding affinity with its natural ligand. The whole sequence of the antigen originates from the SARS-CoV-2 RBD domains of the B.1.1.7 (Alpha) and B.1.351 (Beta) variants, which have been shown by Ramanathan *et al*. to bind ACE2 with increased affinity^34^. We were able to obtain the antigen at high purity, which is consistent with its use as an active drug substance in a vaccine. CHO, the expression system selected to produce this antigen, has been a workhorse to produce monoclonal antibodies and other protein-based therapeutic entities for decades^35^. It has been fully accepted for this purpose by regulatory agencies worldwide.

The PHH-1V vaccine candidate was shown to be safe in mice since it did not cause clinical signs (general and local) nor was there any bodyweight loss attributable to the vaccine composition in either immunised BALB/c or K18-hACE2 mice. Although the histological evaluation of the injection sites revealed mild lesions with cellular infiltrates in a few vaccinated animals, these were attributable to the local innate immune response induced upon injection with adjuvant-containing vaccines. Moreover, we have consistently observed an adequate safety profile in other animal species in which the PHH-1V vaccine candidate has been tested, such as rats, rabbits, cynomolgus monkeys and pigs (manuscript in preparation). The SQBA adjuvant used in this vaccine might be related to the good tolerability shown in these animal models.

Regarding the RBD-binding antibodies humoral response, a dose response was observed on D35/D37 upon vaccination with RBD heterodimer doses of 0.04, 0.2 and 1 μg/dose. However, this response saturates with higher immunisation doses. Significant total IgG titres were observed after just a single dose for doses of 0.2 μg and above. These results suggest a good potency profile for the antigen included in the PHH-1V vaccine candidate, with similar or superior performance to other previously reported immunogens based on similar platforms^36,37^. Moreover, potent pseudovirus-neutralising activity against the Alpha variant was elicited by the 0.2-μg RBD fusion heterodimer/dose immunisation, reaching the highest titres with the 20-μg RBD fusion heterodimer/dose immunisation. Furthermore, a robust pseudovirus-neutralising activity of sera from mice immunised with 20 μg of RBD fusion heterodimer/dose was confirmed against the Beta, Delta and Omicron BA.1 variants. This cross-reactivity was previously confirmed in earlier exploratory trials, where no significant differences were observed in the pseudovirus-neutralising titres against the Alpha, Beta, and Gamma variants in mice upon vaccination with 20-μg RBD fusion heterodimer/dose. Notably, the pseudovirus neutralisation assays from this work were performed in the same laboratory and under identical conditions as those that were previously reported to have a good correlation with live virus neutralisation assays^38^ which highlights the biological relevance of the neutralising antibody titres that were obtained. Even though the recombinant protein RBD fusion heterodimer was designed to elicit a response against the SARS-CoV-2 Alpha and Beta variants, our data also demonstrate a further neutralising activity against the Omicron BA.1 variant, the dominant variant around the world at the time when the study was designed^39,40^. Indeed, our antigen contains several mutations that are key for being considered of high concern, mutations which are present in currently designated VoCs and which could potentially arise in future variants. That includes the E484K substitution present in the Beta and B.1.621 (mu) variants, as well as the K417N and N501Y mutations present in the Omicron BA.1 variant. E484K is related to immune evasion and reduced antibody neutralisation, compromising the efficacy of the original approved vaccines^41^.

Regarding cellular response upon vaccination, the ICS data indicate that immunisation with the highest dose of 20-μg RBD fusion heterodimer/dose induced a robust Th1-dominant response with activation of CD4^+^ and CD8^+^ T cells expressing IFN-γ and IL-2. Notably, ICS detected no significant IL-4 expression in the splenocytes from immunised animals. However, IL-4 ELISpot assays detected the expression of this Th2 cytokine in splenocytes from both immunised groups. Specifically, according to the ELISpot results, immunisation with the 5-μg RBD fusion heterodimer/dose elicited a Th2-biased response, while immunisation with the 20-μg RBD fusion heterodimer promoted a balanced Th1/Th2 response. These differences in the cytokine expression between both assays might be explained by the differences in the incubation time of the splenocytes after the RBD peptide pool stimulation, which was 48 h for the ELISpot compared to 5 h for the ICS. Furthermore, the experimental conditions (number of splenocytes and incubation time) assayed to detect IFN-γ and IL-4 via ELISpot were different; hence, these data must be interpreted carefully. Th1 immunity is known to be protective for most infections since it promotes humoral immunity as well as phagocytic and cytotoxic T cell activity, whereas the Th2 response assists with the resolution of inflammation^42^. Based on the ICS data, immunisation with the 20-μg RBD fusion heterodimer/dose seems to induce a polarised Th1 immune response.

Extracellular cytokine production was also measured by Luminex Multiplex in supernatants from splenocytes after 48 h of stimulation, where a balanced production of Th1 (TNF-α, IL-2) and Th2 (IL-5 but no IL-4 nor IL-6) cytokines was found in vaccinated mice. Notably, IFN-γ was not detected by Luminex, probably due to the early expression of this factor and its rapid degradation. Importantly, IL-10 was not detected in the supernatants, which indicates that the immunisation with PHH-1V did not elicit an anti-inflammatory response after the restimulation of splenocytes with RBD peptide pools.

The IgG2a/IgG1 ratio was measured to assess Th1/Th2 polarization after the prime-boost immunisation. IgG1 is produced during any type of immune response, while IgG2a is mainly produced during a Th1-polarised immune response^43^. Mice immunised with either the 5-μg or the 20-μg RBD fusion heterodimer/dose induced RBD-binding antibodies of both IgG2a and IgG1 subclasses, with an IgG2a/IgG1 ratio near 0.8, indicating a balanced Th1/Th2 response upon PHH-1V vaccination.

Thus, all the data suggest that PHH-1V immunisation with the 20-μg RBD fusion heterodimer/dose elicits a robust CD4^+^ and CD8^+^ T cell response with an early expression of Th1 cytokines upon restimulation *in vitro*, and balanced Th1/Th2 cytokine production after 48 h post-stimulation.

Regarding the preclinical efficacy of the PHH-1V vaccine candidate, it was tested at 2 different doses, 10 μg and 20 μg of RBD fusion heterodimer/dose, in K18-hACE2 mice. Upon the SARS-CoV-2 (Wuhan/D614G strain) challenge, vaccinated animals were able to overcome the infection since neither clinical signs nor bodyweight loss were detected. By contrast, all non-vaccinated and infected animals reached the endpoint criteria on D5 or D6 post-challenge and had to be euthanised. Furthermore, this group of animals experienced remarkable weight loss from D3 post-challenge onwards due to the SARS-CoV-2 infection. Therefore, our data show 100% efficacy in preventing mortality and bodyweight loss in infected K18-hACE2 mice upon PHH-1V vaccination.

In addition, immunisation with either the 10-μg or the 20-μg RBD fusion heterodimer/dose of PHH-1V reduced the viral load measured via qPCR in the lungs, nasal turbinate, and brain in K18-hACE2 mice. The viral load excretion measured in oropharyngeal swabs was also reduced upon challenge in vaccinated animals. Moreover, differences in the viral load after the SARS-CoV-2 challenge between vaccinated animals and infected non-vaccinated control animals were also found in other respiratory (trachea and pharynx) and systemic (spleen and heart) organs. Notably, when RT-qPCR positive samples were titrated to determine the infective viral load, most of the samples of vaccinated animals showed negative results, whereas most of the samples of the infected control group resulted in significantly higher viral loads. Taken together, these results suggest less viral replication in vaccinated mice, which discards antibody-dependent enhancement (ADE) of the infection upon vaccination. Indeed, RBD is known to pose a low potential for risk of ADE because antibodies against this domain block receptor binding ^44^. Likewise, the histopathological evaluation of tissues from vaccinated mice showed no lesions in the brain and mild lesions in the lungs upon SARS-CoV-2 infection. By contrast, infected control mice displayed moderate lesions in the lungs and brain, which is consistent with the high viral loads detected in this group.

Notably, preclinical efficacy of PHH-1V vaccine was also assessed in K18-hACE2 mice against Beta, Delta and Omicron BA.1 VoCs. Overall, clinical signs were observed in all non-vaccinated animals infected with the Delta variant, and also in two non-vaccinated animals infected with the Beta variant, while no clinical signs were found in animals vaccinated with 20-μg RBD fusion heterodimer/dose (except 1 mouse). Furthermore, vaccination with 20-μg RBD fusion heterodimer/dose significantly reduced viral load in lungs and oropharyngeal swabs from animals challenged with the Beta and Delta variants. Histopathological scores were also higher in non-vaccinated animals infected with the Beta or Delta variants compared to PHH-1V vaccinated animals. Although there were no significant differences in survival curves and no major clinical signs in the different groups challenged with the Omicron BA.1 variant, viral load was also reduced in lungs from PHH-1V vaccinated animals compared to infected control animals. The reduction of infectivity and pathogenesis of the Omicron BA.1 variant in K18-hACE2 mice has been reported previously in various studies, including a study in which mRNA-1273 protective efficacy was evaluated against Omicron BA.1^45,46^. Hence, PHH-1V vaccination can reduce and control infection of different VoCs, including Omicron BA.1, in lower respiratory airways. This is critical to mitigate the current pandemic situation, although further studies will have to confirm our findings in human subjects.

Overall, in this study, the PHH-1V vaccine has been shown to be safe and immunogenic in mice, inducing RBD-binding and neutralising antibodies. Mice immunised with 20 μg of recombinant protein RBD fusion heterodimer/dose showed neutralising activity against the Alpha, Beta, Delta and Omicron BA.1 variants. Likewise, immunisation with 20 μg of RBD fusion heterodimer/dose elicited robust activation of CD4^+^ and CD8^+^ T cells, producing an early Th1 response upon *in vitro* restimulation. Importantly, vaccination with either the 10-μg or the 20- μg RBD fusion heterodimer/dose prevented weight loss and clinical signs (including mortality) upon SARS-CoV-2 challenge in mice. Both tested doses reduced viral loads in several organs and prevented the infective viral load in the lungs and brain upon experimental infection. In addition, immunisation with 20 μg of RBD recombinant protein fusion heterodimer reduced the infective viral load in the upper respiratory tract (nasal turbinate). Most importantly, vaccination with 20 μg of RBD recombinant protein fusion heterodimer also reduced the infective viral load of Beta, Delta and Omicron BA.1 variants in the lower respiratory tract. Besides the efficacy and safety features of PHH-1V, this second-generation COVID-19 vaccine is easy to adapt to potential emergent SARS-CoV-2 variants, allowing for the inclusion of up to 2 different RBDs to generate cross-immunity against emergent variants. The PHH-1V vaccine candidate showed promising preclinical data and is currently being evaluated in Phase I/lla (NCT05007509), Phase llb (NCT05142553) and Phase III (NCT05246137) clinical trials^47^.

## 4. Limitations of study

Although the PHH-1V vaccination reduced the infective viral load of Omicron BA.1 variant in the lower respiratory tract, the K18-hACE2 mice model has shown to be asymptomatic upon the SARS-CoV-2 Omicron BA-1 experimental infection. Therefore other animal model should be addressed to evaluate the prevention of clinical signs after an experimental infection with the SARS-CoV-2 Omicron BA-1 variant.

PHH-1V vaccination reduced the infective viral load in the upper respiratory tract after an experimental infection with SARS-CoV-2. However, the local immune response in the upper respiratory tract needs to be analysed in further studies.

## 5. Acknowledgements

We would like to thank Marta Ribó, Ariadna Pararols, Rubén Hernández and Helena Sánchez for their assistance in animal care; Jèssica Gómez and Maria Bosch for their support in the tabulation of raw data; Anna Moya, Mireia Muntada, Lidia Galindo and Núria Elias for the ELISA analysis and protein characterisation; Marta Bau for her assistance in the production of the vaccine antigen; Claudia Millán for helping with the modelling and visualisation of the heterodimeric RBD structure; David Solanes Foz for his support to the ABSL3 Unit; the CMCiB Ethical Committee members (Jorge Carrillo, Eulália Genescà, Carol Galvez, Carol Soler and Patricia Fachal) for their support in the ethical review of animal procedures; the IGTP Biosafety Committee members (Pere-Joan Cardona, Juliá Blanco, Meritxell Carrió, Natalia Ruiz, David Izquierdo and Noemí Parraga) for their support in the review of biosafety procedures; Glòria Pujol and Eduard Fossas for their assistance in the review of the manuscript; Francisco Díaz-Sáez and Anna Muñoz from Alta Medical Services for providing medical writing support; and Adrián Lázaro-Frías from Dynamic Science S.L.U. (Evidenze Clinical Research) for providing medical writing support. We gratefully acknowledge editorial support from Sarah Marshall. This project was partially funded by the Centre for the Development of Industrial Technology (CDTI, IDI20210115), a public organisation answering to the Spanish Ministry of Science and Innovation. Javier Iglesias-Fernández is supported by the Torres Quevedo Programme grant no. PTQ2020-011291 by the Spanish Ministry of Science and Innovation.

## 6. Author Contributions

Conceptualisation, Investigation, Methodology, Resources, Formal Analysis and Data Curation: A.B., A.P., L.F., L.M., J.C., A.F., A.Mo., M.C., M.M., G.B., E.P.M., M.R., R.M., L G., E.P.M., E.P.V., J.P., C.P., T.P.P. & A.M.; Writing Original Draft, Writing – Review & Editing, Visualisation and Supervision: A.B. & A.P.; Investigation: J.R., N.R., M.P., C.L.-O., J.V-A., L.F.-B., J.S., E.P., S.M., B.T., R.O., P.C.R., M.S.-O., J.D., R.A., Y.R. & J.L-C.; Resources: J.l-F, J.V.-A., C.L.-O., J.S., B.T., B.C., J.B., A.M., J.D., Y.R. & S.C.; Writing – Review & Editing: A.Mo, J.V.-A., C.L.-O., J.S., E.P., B.T., B.C., J.B., A.M., J.D., S.C., J.G-P & J.B.; Supervision: J.V.-A., J.S., B.C., J.B., A.M., J.G.-P. & J.B.; Project Administration: L.F.; Conceptualisation: T.P.C. & C.G.

## 9. STAR Methods

### RESOURCE AVAILABILITY

#### Lead contact

Requests for further information or data should be directed to and will be fulfilled by the lead contact, Antoni Prenafeta.

#### Materials availability

Project-related biological samples are not available since they may be required by regulatory agencies or by HIPRA during the clinical development of the vaccine.

#### Data and code availability

- Data reported in this study cannot be deposited in a public repository because the vaccine is under clinical evaluation. Upon request, and subject to review, the lead contact will provide the data that support the reported findings.
- This paper does not report original code.
- Any additional information required to reanalyse the data reported in this paper is available from the lead contact upon request.

### EXPERIMENTAL MODEL AND SUBJECT DETAILS

#### Animals

BALB/c mice (Envigo, #162) and B6.Cg-Tg(K18-ACE2)2Prlmn/J (K-18-hACE2) transgenic mice (Jackson Laboratories, #034860) were used as animal models. All procedures that involved BALB/c mice were conducted in accordance with the European Union Guidelines for Animal Welfare (Directive 2010/63/EU) and approved by the Ethics Committee of HIPRA Scientific S.L.U. and the Department of Territory and Sustainability of the Catalan Government (file: 11388). The experimental procedure that involved the use of K18-hACE2 mice was conducted in accordance with the European Union Guidelines for Animal Welfare (Directive 2010/63/EU) and was approved by the CMCiB Ethics Committee and the Department of Territory and Sustainability of the Catalan Government (file: 11490). The animal study design followed the principles of the 3Rs and animal welfare.

Forty-eight 5-week-old female BALB/c mice were allocated to 6 groups (n=8) and were used for safety and immunogenicity assays. BALB/c mice were injected intramuscularly with a 0.1 mL/dose of the test vaccine, distributed equally in both hind legs (2 x 50 μL), on days 0 (prime) and 21 (boost). Group A was vaccinated with PBS; group B was immunised with the 0.04-μg recombinant protein RBD fusion heterodimer/dose; group C was immunised with the 0.2-μg recombinant protein RBD fusion heterodimer/dose; group D was immunised with the 1-μg recombinant protein RBD fusion heterodimer/dose; group E was immunised with the 5-μg recombinant protein RBD fusion heterodimer/dose; and group F was immunised with the 20- μg recombinant protein RBD fusion heterodimer/dose. These animals were monitored daily for clinical signs and bodyweight was recorded weeKly until D35/D37; at that time, the animals were euthanised and tissues were collected. Animals were watered and fed *ad libitum* with Premium Scientific Diet SAFE*^®^* A04 (Safe-lab). Animals were kept on Arbocel^®^ small functional cellulose pellets (Rettenmaier Ibérica, S. L.) with a light/dark cycle of 12 h at a 22 ºC ±2 ºC in optimum hygienic SPF conditions behind a barrier system under positive pressure with 37 air room renovations per hour. The animals were housed in a stainless-steel rack with polycarbonate cages (530 x 280 x 150 mm) with stainless steel covers equipped with environmental enrichment (nest material: cellulose paper and wood-wool, one PET roll and a PET plastic enrichment dome and a red translucent wheel). The animals were identified with a cage card and individual fur dye. A precision scale (Sartorius, model 112, 6.1 kg with 0.01 g resolution) was used to record the animals’ weights.

For further safety, immunogenicity, and efficacy assays, sixty-two (31F + 31M) 4/5-week-old K18-humanised ACE2 (hACE2) mice were allocated to 4 groups (n=18; 9F + 9M, except for the placebo group: n=8; 4F + 4M). Specifically, group A was intramuscularly injected with PBS and non-infected; group B was injected with PBS and infected with SARS-CoV-2; group C was vaccinated with 10 μg/dose of recombinant protein RBD fusion heterodimer and infected with SARS-CoV-2; and group D was vaccinated with 20 μg/dose of recombinant protein RBD fusion heterodimer and infected with SARS-CoV-2. Animals from satellite subgroups were euthanised on D35 to assess the immunological response of the vaccinated group. Challenged animals were chronologically euthanised on D37, D39 and D42 (males)/D43 (females). Several tissue samples were collected for further analyses. Efficacy were also assessed against Beta, Delta and Omicron BA.1 variants in three different studies using forty-two 4/5-week-old K18-humanised ACE2 (hACE2) mice. Each study had 2 groups of 18 animal (9F + 9M) and a placebo group of 6 animals (3F + 3M). In particular, group A was intramuscularly injected with PBS and non-infected; group B was injected with PBS and infected with Beta, Delta or Omicron BA.1 SARS-CoV-2 variant; and group C was vaccinated with 20 μg/dose of recombinant protein RBD fusion heterodimer and infected with Beta, Delta or Omicron BA.1 SARS-CoV-2 variant. Animals from satellite subgroups were euthanised on D35 to assess the immunological response of the vaccinated group. Challenged animals were also chronologically euthanised on D37, D39 and D42 (males)/D43 (females) in order to collect several tissue samples for RT-qPCR, virus titration and histopathology. Animals were watered and fed *ad libitum* with TeKlad Global 16% Protein Rodent Diet (Envigo, #2916). Animals were kept on cellulose pellets from Rettenmaier Ibérica. Animals were housed in a ventilated rack, model Blue Line Next/Boxunss (Tecniplast, #1145T00SUV-CP), equipped with environmental enrichment (nest material: cellulose paper and cardboard roll). Animals were kept with a light/dark cycle of 12h at 22 ºC ± 2 ºC, with negative room pressure and 20 air renovations per hour. The animals were identified with a dorsal subcutaneous microchip (Trovan, #ID100-B 1.4 Mini transponder). A reader with an integrated scale (Trovan, model 2812005) was used to record the animals’ weights. Cibertec was used as the anaesthesia equipment.

In order to comply with animal welfare regulations, K18-hACE mice were injected with a 0.1 mL/dose of the test vaccine, distributed equally in both hind legs (2 x 50 μL). Vaccines were injected intramuscularly following a two-dose prime-and-boost schedule: 1st dose (prime) on D0 and 2nd dose (boost) on D21. Animals from satellite subgroups were euthanised on D35 to assess the immunological response of the vaccinated group. The SARS-CoV-2 challenge was performed through intranasal inoculation with the strain SARS-CoV-2 Catalonia 02 on a subset of animals on D35 with 25 μL in each nostril (10^3^ TCID_50_/mice in 50 μL/mice). This strain (GISAID ID: EPI_ISL_471472), which included the following mutations compared to Wuhan strain, D614G (Spike), K837N (NSP3), P323L (NSP12), was isolated from a male patient from Barcelona, who showed respiratory symptoms. The intranasal experimental infection was performed under sedation with isoflurane 4-5%. The same procedures were followed for infections with SARS-CoV-2 Beta (B.1.351; GISAID EPI _ISL_1663571), Delta (B.1.617.2; GISAID EPI_ISL_3342900) and Omicron (BA.1; GISAID EPI_ISL_8151031) variants.

BALB/c mice vaccination and sampling were performed at HIPRA (Girona, Spain). K18-hACE2 mice vaccination, SARS-CoV-2 challenge, and sampling were performed in the ABSL3 unit of the Comparative Medicine and Bioimage Centre of Catalonia of the Germans Trias i Pujol Research Institute (Badalona, Spain). The protocol followed is depicted in **Figure 2**.

#### Cell lines

HEK293T cells overexpressing WT human ACE-2 (Integral Molecular, USA) were used as target in the pseudovirus-based neutralisation assay. Cells were maintained in T75 flasks with Dulbecco⍰s Modified Eagle⍰s Medium (DMEM) supplemented with 10% FBS and 1 μg/mL of Puromycin (Thermo Fisher Scientific, USA).

Expi293F cells (Thermo Fisher Scientific) are a HEK293 cell derivative adapted for suspension culture, which were used for SARS-CoV-2 pseudovirus production. Cells were maintained under continuous shaking in Erlenmeyer flasks following the manufacturer’s guidelines.

Vero E6 (ATCC CRL-1586) cell monolayers were cultured for 3 days at 37 ºC and 5% CO2 in DMEM (GIBCO) supplemented with 100 U/mL penicillin, 100 μg/mL streptomycin and 2 mM glutamine (all reagents from Thermo Fisher Scientific). CHO cells were cultured in a bioreactor in a chemically defined media, at 36-38 ºC with a pH 6.80 - 7.40, 5-8% CO_2_ for 60-108 hours with stirring (tip speed 0.4-1) and glucose 2-9 g/L.

### METHOD DETAILS

#### Computational modelling of antigen constructs

For the estimation of the protein-protein interaction energies of the two studied construct variants (B.1.351-B.1.1.7 and B.1.1.7-B.1.351), AlphaFold2^33^ models were generated for each system followed by selection of an individual candidate conformation per construct variant, based on the strongest protein-protein interaction energies identified with the pyDock scoring function^48^. Selected candidates were used as starting models for Molecular Dynamics (MD) simulations with the Amber18 software package^49^. Each protein was immersed in a preequilibrated octahedral water box with a 12-Å buffer of TIP3P water molecule^50^ using the leap module, resulting in the addition of ~26,000 solvent molecules. The systems were subsequently neutralised by addition of explicit counterions (Na^+^ and Cl^-^). All calculations were done using the widely tested ff14SB Amber protein force field^51^. A two-stage geometry optimisation approach was performed, consisting of an initial minimisation of solvent molecules and ions (imposing protein restraints of 500 kcal·mol^-1^Å-2) followed by an unrestrained minimisation of all atoms in the simulation cell. The systems were then gently heated using six 50-ps steps, incrementing the temperature 50 K each step (0–300 K) under constant volume and periodic boundary conditions. Next, both systems were then equilibrated without restraints for 2 ns at a constant pressure of 1 atm and temperature of 300 K. Finally, 500 ns MD production simulations were performed for each of the systems in the NVT ensemble and periodic boundary conditions. Models of both constructs bound to a hACE2 dimeric receptor were manually built based on available x-ray crystal structures (PDB code: 6M17) and MD parameterised following the protocol described above. Production runs of 100 ns were calculated for each system studied.

The RBD-RBD interaction energies between candidate constructs and construct-hACE2 receptor were calculated using the MM-GBSA method in Amber18^52^. For each PHH-1V construct, the MM-GBSA calculation was performed using 300 snapshots over the last 300 ns of the simulation with 1 ns interval with the MMPBSA.py module in Amber 18 with an ionic strength equal to 0.1 M.

#### Recombinant RBD fusion heterodimer characterisation

The antigen was produced in a bioreactor based on a selected stable CHO clone. A fed-batch strategy was used for high-cell-density cultivation and expression of the RBD fusion heterodimer. Upon harvest, the cell broth was clarified by depth filtration. The clarified supernatant was further purified via sequential chromatography. The purified antigen was then buffer exchanged by tangential flow filtration and filter sterilised. Purity and integrity were evaluated by SDS-PAGE with Bolt^™^ 4 to 12% Bis-Tris gels (Thermo Fisher, ref. NW04120BOX), stained with One-Step Blue Protein Gel Stain (Biotium, ref. 21003), and by SEC-HPLC with an Xbridge Protein BEH SEC (Waters, ref. 186009160) connected to an HP1100 system (Agilent Technologies).

The affinity test of the RBD heterodimer with human ACE2 by surface plasmon resonance (SPR) was performed by ACROBiosystems. The Fc-tagged ACE2 (AC2-H5257, ACROBiosystems) was immobilised in a Series S Sensor Chip CM5 (Cytiva) on a Biacore T200 (Cytiva) using the Human Antibody Capture Kit (Cytiva). The affinity measure was obtained using 8 different RBD heterodimer concentrations. The antigen structure simulations were performed with UCSF ChimeraX^53^.

#### SARS-CoV-2 recombinant protein RBD heterodimer adjuvanted vaccines

The purified RBD fusion heterodimer was formulated with the SQBA adjuvant, an oil-in-water emulsion produced by HIPRA. The PHH-1V vaccine was tested at different concentrations: 0.04 μg, 0.2 μg, 1 μg, 5 μg and 20 μg of RBD fusion heterodimer/dose for the safety and immunogenicity assays in BALB/c mice. For efficacy assessment in the K18-hACE2 mice animal model, the vaccine was tested at 10 μg and 20 μg of fusion heterodimer/dose. The placebo vaccines were prepared with phosphate-buffered saline (PBS).

#### Analysis of SARS-CoV-2-specific antibodies

Serum binding antibodies against SARS-CoV-2 RBD were determined by ELISA (HIPRA). MaxiSorp plates (Nunc, Roskilde, Denmark) were coated with 100 ng/well RBD protein (Sino Biologicals, Beijing, China) and blocked with 5% non-fat dry milk (Difco^™^ Skim Milk, BD, FranKlin Lakes, NJ, USA) in PBS. Wells were incubated with serial dilutions of the serum samples and the bound total IgG specific antibodies were detected by peroxidase-conjugated Goat Anti-Mouse IgG (Sigma-Aldrich, St. Louis, MO, USA). Finally, wells were incubated with K-Blue Advanced Substrate (Neogen, Lansing, Ml, USA) and the absorbance at 450 nm was measured using a microplate reader (Versamax microplate reader, Molecular Devices, San Jose, CA, USA). The mean value of the absorbance was calculated for each dilution of the serum sample run in duplicate. Isotypes IgG1 and IgG2a were detected using Peroxidase AffiniPure Goat Anti-Mouse IgG, Fcγ subclass 1 specific, and Peroxidase AffiniPure Goat AntiMouse IgG, Fey subclass 2a specific, (Jackson ImmunoResearch, Cambridge, UK), respectively. The endpoint titre of RBD-specific total IgG binding antibodies was established as the reciprocal of the last serum dilution yielding 3 times the mean optical density of the negative control of the technique (wells without serum added).

#### Pseudovirus neutralisation assay

Neutralising antibodies in serum against SARS-CoV-2 Wuhan (original sequence) and the Alpha, Beta, Gamma, and Delta variants were determined by a pseudoviruses-based neutralisation assay (PBNA) at IRSICaixa (Barcelona, Spain) using an HIV reporter pseudovirus that expresses the S protein of SARS-CoV-2 and luciferase. To generate pseudoviruses, Expi293F cells were transfected using ExpiFectamine293 Reagent (Thermo Fisher Scientific) with pNL4-3.Luc.R-.E- and SARS-CoV-2.SctΔ19 at a 24:1 ratio, respectively^54^. pNL4-3.Luc.R-.E-was obtained from the NIH AIDS Reagent Program and SARS-CoV-2.SctΔ19 was generated by GeneArt from the full protein sequence of SARS-CoV-2 spike with a deletion of the last 19 amino acids in C-terminal,24 human-codon optimized and inserted into pcDNA3.4-TOPO. Control pseudoviruses were obtained by replacing the S protein expression plasmid with a VSV-G protein expression plasmid (pVSV-G) described before^55^. Supernatants were harvested 48 hours after transfection, filtered at 0.45 mm, frozen, and titrated on HEK293T cells overexpressing WT human ACE-2 (Integral Molecular, USA).

For the neutralisation assay, 200 TCID_50_ of pseudovirus supernatant was preincubated with serial dilutions of the heat-inactivated serum samples for 1 h at 37 °C and then added onto ACE2 overexpressing HEK293T cells. After 48 h, cells were lysed with britelite plus luciferase reagent (PerkinElmer, Waltham, MA, USA). Luminescence was measured for 0.2 s with an EnSight multimode plate reader (PerkinElmer). The neutralisation capacity of the serum samples was calculated by comparing the experimental RLU calculated from infected cells treated with each serum to the max RLUs (maximal infectivity calculated from untreated infected cells) and min RLUs (minimal infectivity calculated from uninfected cells) and expressed as the neutralisation percentage: neutralisation^38^ (%) = (RLUmax–RLUexperimental)/(RLUmax–RLUmin) * 100. IC_50_ were calculated by plotting and fitting neutralisation values and the plasma dilution log to a 4-parameters equation in Prism 9.0.2 (GraphPad Software, USA).

#### Intracellular cytokine staining (ICS)

ICS was performed by the Infection Biology Group at the Department of Experimental and Health Sciences, Universitat Pompeu Fabra (DCEXS-UPF, Barcelona, Spain). Spleens from female mice were mechanically disrupted onto a 40-μM cell strainer and incubated in 5 mL of 0.15 M ammonium chloride buffer for 5 min at room temperature (RT) for red blood cell lysis. Cells were then washed in RPMI (Gibco, Tavarnuzze, Italy) supplemented with 10% FBS, 1% penicillin/streptomycin, 0.05 mM-Mercaptoethanol and 1 mM sodium pyruvate (cRPMI). Two million splenocytes per well (96-well plate) were stimulated in vitro under three conditions: (i) a 1:1 mix of the peptide libraries (PepMix^™^) from the B.1.1.7 (Alpha variant) and B.1.351 (Beta variant) lineages covering the entire RBD of the SARS-CoV-2 S protein; (ii) cRPMI (negative control); and (iii) PMA + lonomycin (positive control) for 5 h at 37 ºC 5% CO_2_ in cRPMI in the presence of Brefeldin A (Sigma-Aldrich) for the last 3 h before antibody staining. The final concentrations used were 1 μg/mL of each peptide of the RBD peptide pool, 15 ng/mL of PMA (Sigma-Aldrich) and 250 ng/mL of ionomycin (Sigma-Aldrich). For flow cytometric analysis, equal numbers of cells were stained with Fixable Viability Stain 780 (BD Biosciences, New Jersey, NJ, USA) in PBS for 15 min at RT followed by staining with antibodies against CD3, CD4, CD8 and CD44 for 20 min on ice in FACS buffer (PBS: 5% FCS, 0.5% BSA, 0.07%NaN3). Cells were then fixed for 20 min on ice with 2% formaldehyde and stained with antibodies against intracellular proteins (IFN-γ, TNF-α, IL-2 and IL-4) for 20 min on ice in perm/wash buffer (PBS: 1% FCS, NaN3 0.1%, saponin 0.1%). All antibodies were purchased from either BD Biosciences, Thermo Fisher or BioLegend (see Table S1 for more details). Samples were processed on an Aurora analyser (Cytek, Fremont, CA, USA). FACS data were analysed using FlowJo 10 software (Tree Star Inc., Ashland, OR, USA). The gating strategy followed in the analysis is depicted in **Figure S6**. The stain index was calculated by subtracting the mean fluorescence intensity (MFI) of the unstained or fluorescence minus one (FMO) control from the MFI of the stained samples and dividing it by two times the standard deviation of the unstained population. Background cytokine expression in the no-peptide (cRPMI) condition was subtracted from that measured in the RBD peptide pool for each mouse. To calculate the Th1 response in CD4^+^ and CD8^+^, the Boolean tool of the FlowJo software was used.

#### Mouse cytokine assay

The cytokine assay was performed by the Infection Biology Group at the Department of Experimental and Health Sciences, Universitat Pompeu Fabra (DCEXS-UPF, Barcelona, Spain). Splenocytes from female mice were seeded at 1.1 × 10^6^ cells/well in 24-well plates and stimulated with a 1:1 mix of the RBD overlapping peptides from B.1.1.7 (Alpha variant) and B.1.1351 (Beta variant) lineages (1 μg/mL each). cRPMI media was used as a negative control and PMA (15 ng/mL) + ionomycin (250 ng/mL) as a positive control. The supernatants were harvested after 48 h incubation at 37 °C and a panel that quantifies the cytokines IL-2, IL-4, IL-5, IL-6, IL-10, IFN-γ and TNF-α (Luminex Multiplex, Invitrogen, Waltham, MA, USA) was run according to the manufacturer’s instructions. These measurements were performed at Veterinary Clinical Biochemistry Service, Faculty of Veterinary Medicine, Universitat Autònoma de Barcelona (UAB, Barcelona, Spain).

#### IFN-γ and IL-4 ELISpot assays

ELISpot assays were performed with mouse IFN-γ and IL-4 ELISpot PLUS kits according to the manufacturer’s instructions (3321-4HPT-10 and 3311-4HPW-10, Mabtech, Herndon, VA, USA). A total of 2.5 × 10^5^ or 4 x 10^5^ splenocytes from female mice were seeded per well for the IFN-γ and IL-4 tests, respectively, and ex *vivo* stimulated either with the 1:1 mix of the RBD overlapping peptides from the B.1.1.7 (Alpha variant) and B.1.351 (Beta variant) lineages (1 μg/mL each), or with complete cRPMI (negative control) or with concanavalin A (5 μg/mL (Sigma-Aldrich)) (positive control). Each condition was run in duplicates. After an incubation period of 18-20 h (for IFN-γ) or 48 h (for IL-4), the plates were manipulated according to the manufacturer’s instructions. Spots were counted under a dissection microscope. Frequencies of IFN-γ or IL-4-secreting cells were expressed as the number of responding cells per million splenocytes. The number of spots in unstimulated cultures (negative control) was subtracted from the spot count in RBD-stimulated cultures.

#### SARS-CoV-2 genomic RT-qPCR

Total viral load in respiratory tissue samples was determined by RT-qPCR (CReSA, IRTA-UAB, Barcelona, Spain). Viral RNA was extracted from target organs and swabs samples using the IndiMag pathogen kit (Indical Bioscience) on a Biosprint 96 workstation (QIAGEN) according to the manufacturer’s instructions. The RT-qPCR used to detect viral gRNA was performed with the AgPath-ID One-Step RT-PCR Kit (Life Technologies). In brief, 5 μL of RNA were added to 25 μL reaction containing 12.5 μL of 2 × reaction buffer and 1 μL of 25X RT-PCR Enzyme mix provided with ArrayScript^™^ Reverse Transcriptase and AmpliTaq Gold^®^ DNA Polymerase. RT-qPCR targets a portion of the envelope protein gene (position 26,141–26,253 of GenBank NC_004718). The primers and probes used, and their final concentration, were the following: forward, 5⍰-ACAGGTACGTTAATAGTTAATAGCGT-3⍰ [400 nM]; reverse, 5⍰-ATATTGCAGCAGTACGCACACA-3⍰ [400 nM]; probe, 5⍰-FAM-ACACTAGCCATCCTTA CTGCGCTTCG-TAMRA-3⍰ [200 nM]^56^. Thermal cycling was performed at 55 °C for 10 min for reverse transcription, followed by 95 °C for 3 min, and then 45 cycles of 94 °C for 15 s plus 58 °C for 30 s.

#### Virus titration in Vero E6 cell

Virus titres were determined in RT-qPCR positive samples using a standard TCID_50_ assay in Vero E6 cells at CReSA (IRTA-UAB) ^56^. Briefly, each sample was 10-fold diluted in duplicate, transferred in a 96 well plate with a Vero E6 cells monolayer and incubated at 37 °C and 5% CO_2_. Plates were monitored daily under the light microscope and wells were evaluated for the presence of CPE at 5 dpi. The amount of infectious virus was calculated by determining the TCID_50_ using the Reed-Muench method.

#### Histopathology

Histopathological analyses were performed at CReSA (IRTA-UAB). Lower (lungs) respiratory tract, brain, spleen, trachea, heart and skeletal muscle were fixed in 10% buffered formalin and routinely processed for histopathology. Haematoxylin- and eosin-stained slides were examined under optical microscope. Multifocal broncho-interstitial pneumonia, multifocal lymphoplasmacytic rhinitis and non-suppurative meningoencephalitis were evaluated from lung, nasal turbinate and brain lesions, respectively, according to the following semi-quantitative score: 0 (no lesion); 1 (mild lesion, multifocal distribution and less than 10% of tissue affected); 2 (moderate lesion, multifocal distribution and between 10-40% of tissue affected); and 3 (severe lesion, multifocal to diffuse distribution and more than 40% of tissue affected)^56,57^. A European certified (ECVP) pathologist performed a blinded pathology assessment.

### QUANTIFICATION AND STATISTICAL ANALYSIS

Statistical analyses and plots were generated using R (version 4.0.5) or GraphPad Prism (version 9). Unless otherwise specified, all plots depict individual data points for each animal, along with the sample mean and standard deviation. When required, data was either log_10_^-^ or arcsine-transformed (i.e., log-normal and percentage variables, respectively). The exact number (n) used in each experiment is indicated in the caption below each figure.

When testing the effect of one or two factors, one-way ANOVA or linear models were generally employed. For models involving independent observations, the generalised least squares approximation (GLS implementation in the R package nlme) was used to accommodate potential heteroscedasticity. Conversely, for models involving repeated measures, linear mixed effects models were fitted using the lme implementation in the R package nlme. Unless otherwise specified, time, group and their interaction were included in the models as fixed effects, and the experimental subject was considered a random factor. The corresponding random intercept models were fitted to the data using restricted maximum likelihood. Correlation between longitudinal observations as well as heteroscedasticity were included in the models when required with appropriate variance-covariance structures. On the other hand, data violating the assumption of normality, as well as ordinal variables, were analysed using Mann-Whitney or Kruskal-Wallis tests, depending on the number of groups, segregating by timepoint if needed.

Assumptions were tested graphically (using quantile-quantile and residual plots) for both modelling approaches, and model selection was based on likelihood ratio tests or a priori assumptions. The corresponding estimated marginal means were calculated using the R package emmeans.

For post-hoc pairwise comparisons, appropriate tests were employed depending on the nature of the data and the comparisons to perform, with corrections for multiple testing when required (i.e., Holm’s-Bonferroni correction or multivariate t-distribution adjustment). When comparisons against zero-variance groups (all observations having the same value) needed to be performed, one-sample tests were employed instead.

Finally, survival analysis was performed to test the differences in clinical signs and mortality using Kaplan-Meier estimates and the log-rank test. For analyses involving more than two groups, a priori pairwise contrasts were employed upon significant omnibus log-rank tests to control type I error.

Statistically significant differences between groups are indicated with a line on top of each group: ** *p<0.01; * p<0.05; ^+^ 0.05<p<0.1*.

### KEY RESOURCES TABLE

See attached document.

## 10. Supplemental Information

See attached document.

## References

1. WHO (2021). WHO Coronavirus (COVID-19) Dashboard. https://covid19.who.int/.

2. Mathieu, E., Ritchie, H., Ortiz-Ospina, E., Roser, M., Hasell, J., Appel, C., Giattino, C., and Rodés-Guirao, L. (2021). Author Correction: A global database of COVID-19 vaccinations. Nat Hum Behav 5, 956–959. 10.1038/s41562-021-01160-2.

3. Ritchie, H., Rodés-Guirao, L., Appel, C., Giattino, C., Ortiz-Ospina, E., Hasell, J., Macdonald, B., Beltekian, D., and Roser, M. (2022). Coronavirus Pandemic (COVID-19). https://ourworldindata.org/coronavirus.

4. Günl, F., Mecate-Zambrano, A., Rehländer, S., Hinse, S., Ludwig, S., and Brunotte, L. (2021). Shooting at a Moving Target-Effectiveness and Emerging Challenges for SARS-CoV-2 Vaccine Development. Vaccines (Basel) 9. 10.3390/vaccines9101052.

5. Moyo-Gwete, T., Madzivhandila, M., Makhado, Z., Ayres, F., Mhlanga, D., Oosthuysen, B., Lambson, B.E., Kgagudi, P., Tegally, H., Iranzadeh, A., et al. (2021). SARS-CoV-2 501Y.V2 (B.1.351) elicits cross-reactive neutralizing antibodies. bioRxiv. 10.1101/2021.03.06.434193.

6. Mellet, J., and Pepper, M.S. (2021). A COVID-19 Vaccine: Big Strides Come with Big Challenges. Vaccines (Basel) 9. 10.3390/vaccines9010039.

7. Alanagreh, L., Alzoughool, F., and Atoum, M. (2020). The Human Coronavirus Disease COVID-19: Its Origin, Characteristics, and Insights into Potential Drugs and Its Mechanisms. Pathogens 9. 10.3390/pathogens9050331.

8. Vogel, A.B., Kanevsky, I., Che, Y., Swanson, K.A., Muik, A., Vormehr, M., Kranz, L.M., Walzer, K.C., Hein, S., Güler, A., et al. (2020). A prefusion SARS-CoV-2 spike RNA vaccine is highly immunogenic and prevents lung infection in non-human primates. bioRxiv, 2020.2009.2008.280818. 10.1101/2020.09.08.280818.

9. Zhou, P., Yang, X.L., Wang, X.G., Hu, B., Zhang, L., Zhang, W., Si, H.R., Zhu, Y., Li, B., Huang, C. L., et al. (2020). Addendum: A pneumonia outbreak associated with a new coronavirus of probable bat origin. Nature 588, E6. 10.1038/s41586-020-2951-z.

10. Huang, Y., Yang, C., Xu, X.F., Xu, W., and Liu, S.W. (2020). Structural and functional properties of SARS-CoV-2 spike protein: potential antivirus drug development for COVID-19. Acta Pharmacol Sin 41, 1141–1149. 10.1038/s41401-020-0485-4.

11. Yi, C., Sun, X., Ye, J., Ding, L., Liu, M., Yang, Z., Lu, X., Zhang, Y., Ma, L., Gu, W., et al. (2020). Key residues of the receptor binding motif in the spike protein of SARS-CoV-2 that interact with ACE2 and neutralizing antibodies. Cell Mol Immunol 17, 621–630. 10.1038/s41423-020-0458-z.

12. Cao, Y., Wang, J., Jian, F., Xiao, T., Song, W., Yisimayi, A., Huang, W., Li, Q., Wang, P., An, R., et al. (2022). Omicron escapes the majority of existing SARS-CoV-2 neutralizing antibodies. Nature 602, 657–663. 10.1038/s41586-021-04385-3.

13. Liu, Z., Xu, W., Xia, S., Gu, C., Wang, X., Wang, Q., Zhou, J., Wu, Y., Cai, X., Qu, D., et al. (2020). RBD-Fc-based COVID-19 vaccine candidate induces highly potent SARS-CoV-2 neutralizing antibody response. Signal Transduct Target Ther 5, 282. 10.1038/s41392-020-00402-5.

14. Plante, J.A., Mitchell, B.M., Plante, K.S., Debbink, K., Weaver, S.C., and Menachery, V.D. (2021). The variant gambit: COVID-19’s next move. Cell Host Microbe 29, 508–515. 10.1016/j.chom.2021.02.020.

15. WHO (2022). Tracking SARS-CoV-2 variants. https://www.who.int/activities/tracking-SARS-CoV-2-variants.

16. Dai, L., and Gao, G.F. (2021). Viral targets for vaccines against COVID-19. Nat Rev Immunol 21, 73–82. 10.1038/s41577-020-00480-0.

17. Pallesen, J., Wang, N., Corbett, K.S., Wrapp, D., Kirchdoerfer, R.N., Turner, H.L., Cottrell, C.A., Becker, M.M., Wang, L., Shi, W., et al. (2017). Immunogenicity and structures of a rationally designed prefusion MERS-CoV spike antigen. Proc Natl Acad Sci U S A 114, E7348–e7357. 10.1073/pnas.1707304114.

18. EMA (2021). Comirnaty EPAR - Product information. https://www.ema.europa.eu/en/documents/product-information/comirnaty-epar-product-information_en-0.pdf.

19. EMA (2021). Spikevax (previously COVID-19 Vaccine Moderna): EPAR - Product information. https://www.ema.europa.eu/en/documents/product-information/spikevax-previously-covid-19-vaccine-moderna-epar-product-information_en.pdf.

20. EMA (2022). Jcovden (previously COVID-19 Vaccine Janssen). https://www.ema.europa.eu/en/medicines/human/EPAR/icovden-previously-covid-19-vaccine-ianssen.

21. Keech, C., Albert, G., Cho, I., Robertson, A., Reed, P., Neal, S., Plested, J.S., Zhu, M., Cloney-Clark, S., Zhou, H., et al. (2020). Phase 1-2 Trial of a SARS-CoV-2 Recombinant Spike Protein Nanoparticle Vaccine. N Engl J Med 383, 2320–2332. 10.1056/NEJMoa2026920.

22. Francica, J.R., Flynn, B.J., Foulds, K.E., Noe, A.T., Werner, A.P., Moore, I.N., Gagne, M., Johnston, T.S., Tucker, C., Davis, R.L., et al. (2021). Vaccination with SARS-CoV-2 Spike Protein and AS03 Adjuvant Induces Rapid Anamnestic Antibodies in the Lung and Protects Against Virus Challenge in Nonhuman Primates. bioRxiv. 10.1101/2021.03.02.433390.

23. Goepfert, P.A., Fu, B., Chabanon, A.L., Bonaparte, M.I., Davis, M.G., Essink, B.J., Frank, I., Haney, O., Janosczyk, H., Keefer, M.C., et al. (2021). Safety and immunogenicity of SARS-CoV-2 recombinant protein vaccine formulations in healthy adults: interim results of a randomised, placebo-controlled, phase 1-2, dose-ranging study. Lancet Infect Dis 21, 1257–1270. 10.1016/s1473-3099(21)00147-x.

24. Kyriakidis, N.C., López-Cortés, A., González, E.V., Grimaldos, A.B., and Prado, E.O. (2021). SARS-CoV-2 vaccines strategies: a comprehensive review of phase 3 candidates. NPJ Vaccines 6, 28. 10.1038/s41541-021-00292-w.

25. Kleanthous, H., Silverman, J.M., Makar, K.W., Yoon, I.K., Jackson, N., and Vaughn, D.W. (2021). Scientific rationale for developing potent RBD-based vaccines targeting COVID-19. NPJ Vaccines 6, 128. 10.1038/s41541-021-00393-6.

26. Gavi Alliance, C.c.p. (2020). What are protein subunit vaccines and how could they be used against COVID-19? https://www.gavi.org/vaccineswork.

27. Piccoli, L., Park, Y.J., Tortorici, M.A., Czudnochowski, N., Walls, A.C., Beltramello, M., Silacci-Fregni, C., Pinto, D., Rosen, L.E., Bowen, J.E., et al. (2020). Mapping Neutralizing and Immunodominant Sites on the SARS-CoV-2 Spike Receptor-Binding Domain by Structure-Guided High-Resolution Serology. Cell 183, 1024–1042.e1021. 10.1016/j.cell.2020.09.037.

28. Wang, H., Wu, X., Zhang, X., Hou, X., Liang, T., Wang, D., Teng, F., Dai, J., Duan, H., Guo, S., et al. (2020). SARS-CoV-2 Proteome Microarray for Mapping COVID-19 Antibody Interactions at Amino Acid Resolution. ACS Cent Sci 6, 2238–2249. 10.1021/acscentsci.0c00742.

29. Greaney, A.J., Loes, A.N., Crawford, K.H.D., Starr, T.N., Malone, K.D., Chu, H.Y., and Bloom, J.D. (2021). Comprehensive mapping of mutations in the SARS-CoV-2 receptor-binding domain that affect recognition by polyclonal human plasma antibodies. Cell Host Microbe 29, 463–476.e466. 10.1016/j.chom.2021.02.003.

30. Greaney, A.J., Loes, A.N., Gentles, L.E., Crawford, K.H.D., Starr, T.N., Malone, K.D., Chu, H.Y., and Bloom, J.D. (2021). Antibodies elicited by mRNA-1273 vaccination bind more broadly to the receptor binding domain than do those from SARS-CoV-2 infection. Sci Transl Med 13. 10.1126/scitranslmed.abi9915.

31. Lee, W.S., Wheatley, A.K., Kent, S.J., and DeKosky, B.J. (2020). Antibody-dependent enhancement and SARS-CoV-2 vaccines and therapies. Nat Microbiol 5, 1185–1191. 10.1038/s41564-020-00789-5.

32. Cox, J.C., and Coulter, A.R. (1997). Adjuvants--a classification and review of their modes of action. Vaccine 15, 248–256. 10.1016/s0264-410x(96)00183-1.

33. Jumper, J., Evans, R., Pritzel, A., Green, T., Figurnov, M., Ronneberger, O., Tunyasuvunakool, K., Bates, R., Žídek, A., Potapenko, A., et al. (2021). Highly accurate protein structure prediction with AlphaFold. Nature 596, 583–589. 10.1038/s41586-021-03819-2.

34. Ramanathan, M., Ferguson, I.D., Miao, W., and Khavari, P.A. (2021). SARS-CoV-2 B.1.1.7 and B.1.351 Spike variants bind human ACE2 with increased affinity. bioRxiv. 10.1101/2021.02.22.432359.

35. Sharker, S.M., and Rahman, A. (2021). A Review on the Current Methods of Chinese Hamster Ovary (CHO) Cells Cultivation for the Production of Therapeutic Protein. Curr Drug Discov Technol 18, 354–364. 10.2174/1570163817666200312102137.

36. Kuo, T.Y., Lin, M.Y., Coffman, R.L., Campbell, J.D., Traquina, P., Lin, Y.J., Liu, L.T., Cheng, J., Wu, Y.C., Wu, C.C., et al. (2020). Development of CpG-adjuvanted stable prefusion SARS-CoV-2 spike antigen as a subunit vaccine against COVID-19. Sci Rep 10, 20085. 10.1038/s41598-020-77077-z.

37. Tian, J.H., Patel, N., Haupt, R., Zhou, H., Weston, S., Hammond, H., Logue, J., Portnoff, A.D., Norton, J., Guebre-Xabier, M., et al. (2021). SARS-CoV-2 spike glycoprotein vaccine candidate NVX-CoV2373 immunogenicity in baboons and protection in mice. Nat Commun 12, 372. 10.1038/s41467-020-20653-8.

38. Trinité, B., Tarrés-Freixas, F., Rodon, J., Pradenas, E., Urrea, V., Marfil, S., Rodríguez de la Concepción, M.L., Ávila-Nieto, C., Aguilar-Gurrieri, C., Barajas, A., et al. (2021). SARS-CoV-2 infection elicits a rapid neutralizing antibody response that correlates with disease severity. Sci Rep 11, 2608. 10.1038/s41598-021-81862-9.

39. Hadfield, J., Megill, C., Bell, S.M., Huddleston, J., Potter, B., Callender, C., Sagulenko, P., Bedford, T., and Neher, R.A. (2018). Nextstrain: real-time tracking of pathogen evolution. Bioinformatics 34, 4121–4123. 10.1093/bioinformatics/bty407.

40. Nextstrain (2021). Genomic epidemiology of novel coronavirus - Global subsampling. https://nextstrain.org/ncov/gisaid/global/6m.

41. Jangra, S., Ye, C., Rathnasinghe, R., Stadlbauer, D., Krammer, F., Simon, V., Martinez-Sobrido, L., García-Sastre, A., and Schotsaert, M. (2021). SARS-CoV-2 spike E484K mutation reduces antibody neutralisation. Lancet Microbe 2, e283–e284. 10.1016/s2666-5247(21)00068-9.

42. Spellberg, B., and Edwards, J.E., Jr. (2001). Type 1/Type 2 immunity in infectious diseases. Clin Infect Dis 32, 76–102. 10.1086/317537.

43. Snapper, C.M., and Paul, W.E. (1987). Interferon-gamma and B cell stimulatory factor-1 reciprocally regulate Ig isotype production. Science 236, 944–947. 10.1126/science.3107127.

44. An, Y., Li, S., Jin, X., Han, J.B., Xu, K., Xu, S., Han, Y., Liu, C., Zheng, T., Liu, M., et al. (2022). A tandem-repeat dimeric RBD protein-based covid-19 vaccine zf2001 protects mice and nonhuman primates. Emerg Microbes Infect 11, 1058–1071. 10.1080/22221751.2022.2056524.

45. Tarrés-Freixas, F., Trinité, B., Pons-Grífols, A., Romero-Durana, M., Riveira-Muñoz, E., Ávila-Nieto, C., Pérez, M., Garcia-Vidal, E., Perez-Zsolt, D., Muñoz-Basagoiti, J., et al. (2022). Heterogeneous Infectivity and Pathogenesis of SARS-CoV-2 Variants Beta, Delta and Omicron in Transgenic K18-hACE2 and Wildtype Mice. Front Microbiol 13, 840757. 10.3389/fmicb.2022.840757.

46. Ying, B., Scheaffer, S.M., Whitener, B., Liang, C.Y., Dmytrenko, O., Mackin, S., Wu, K., Lee, D., Avena, L.E., Chong, Z., et al. (2022). Boosting with variant-matched or historical mRNA vaccines protects against Omicron infection in mice. Cell 185, 1572–1587.e1511. 10.1016/j.cell.2022.03.037.

47. U.S. National Library of Medicine (2021). Safety and Immunogenicity Study of Recombinant Protein RBD Candidate Vaccine Against SARS-CoV-2 in Adult Healthy Volunteers (COVID-19). Identifier: NCT05007509.

48. Cheng, T.M., Blundell, T.L., and Fernandez-Recio, J. (2007). pyDock: electrostatics and desolvation for effective scoring of rigid-body protein-protein docking. Proteins 68, 503–515. 10.1002/prot.21419.

49. Case, D.A., Ben-Shalom, I.Y., Brozell, S.R., Cerutti, D.S., Cheatham III, T.E., Cruzeiro, V.W.D., Darden, T.A., Duke, R.E., Ghoreishi, D., Gilson, M.K., et al. (2018). AMBER 2018, University of California, San Francisco.

50. Jorgensen, W.L., Chandrasekhar, J., Madura, J.D., Impey, R.W., and Klein, M.L. (1983). Comparison of simple potential functions for simulating liquid water. The Journal of Chemical Physics 79, 926–935. 10.1063/1.445869.

51. Maier, J.A., Martinez, C., Kasavajhala, K., Wickstrom, L., Hauser, K.E., and Simmerling, C. (2015). ff14SB: Improving the Accuracy of Protein Side Chain and Backbone Parameters from ff99SB. J Chem Theory Comput 11, 3696–3713. 10.1021/acs.jctc.5b00255.

52. Kollman, P.A., Massova, I., Reyes, C., Kuhn, B., Huo, S., Chong, L., Lee, M., Lee, T., Duan, Y., Wang, W., et al. (2000). Calculating structures and free energies of complex molecules: combining molecular mechanics and continuum models. Acc Chem Res 33, 889–897. 10.1021/ar000033j.

53. Pettersen, E.F., Goddard, T.D., Huang, C.C., Meng, E.C., Couch, G.S., Croll, T.I., Morris, J.H., and Ferrin, T.E. (2021). UCSF ChimeraX: Structure visualization for researchers, educators, and developers. Protein Sci 30, 70–82. 10.1002/pro.3943.

54. Pradenas, E., Trinité, B., Urrea, V., Marfil, S., Ávila-Nieto, C., Rodríguez de la Concepción, M.L., Tarrés-Freixas, F., Pérez-Yanes, S., Rovirosa, C., Ainsua-Enrich, E., et al. (2021). Stable neutralizing antibody levels 6 months after mild and severe COVID-19 episodes. Med (N Y) 2, 313–320.e314. 10.1016/j.medj.2021.01.005.

55. Sánchez-Palomino, S., Massanella, M., Carrillo, J., García, A., García, F., González, N., Merino, A., Alcamí, J., Bofill, M., Yuste, E., et al. (2011). A cell-to-cell HIV transfer assay identifies humoral responses with broad neutralization activity. Vaccine 29, 5250–5259. 10.1016/j.vaccine.2011.05.016.

56. Brustolin, M., Rodon, J., Rodríguez de la Concepción, M.L., Ávila-Nieto, C., Cantero, G., Pérez, M., Te, N., Noguera-Julián, M., Guallar, V., Valencia, A., et al. (2021). Protection against reinfection with D614- or G614-SARS-CoV-2 isolates in golden Syrian hamster. Emerg Microbes Infect 10, 797–809. 10.1080/22221751.2021.1913974.

57. Vidal, E., López-Figueroa, C., Rodon, J., Pérez, M., Brustolin, M., Cantero, G., Guallar, V., Izquierdo-Useros, N., Carrillo, J., Blanco, J., et al. (2022). Chronological brain lesions after SARS-CoV-2 infection in hACE2-transgenic mice. Vet Pathol 59, 613–626. 10.1177/03009858211066841.

