## Supplementary material for "Preclinical evaluation of a COVID-19 vaccine candidate based on a recombinant RBD fusion heterodimer of SARS-CoV-2"

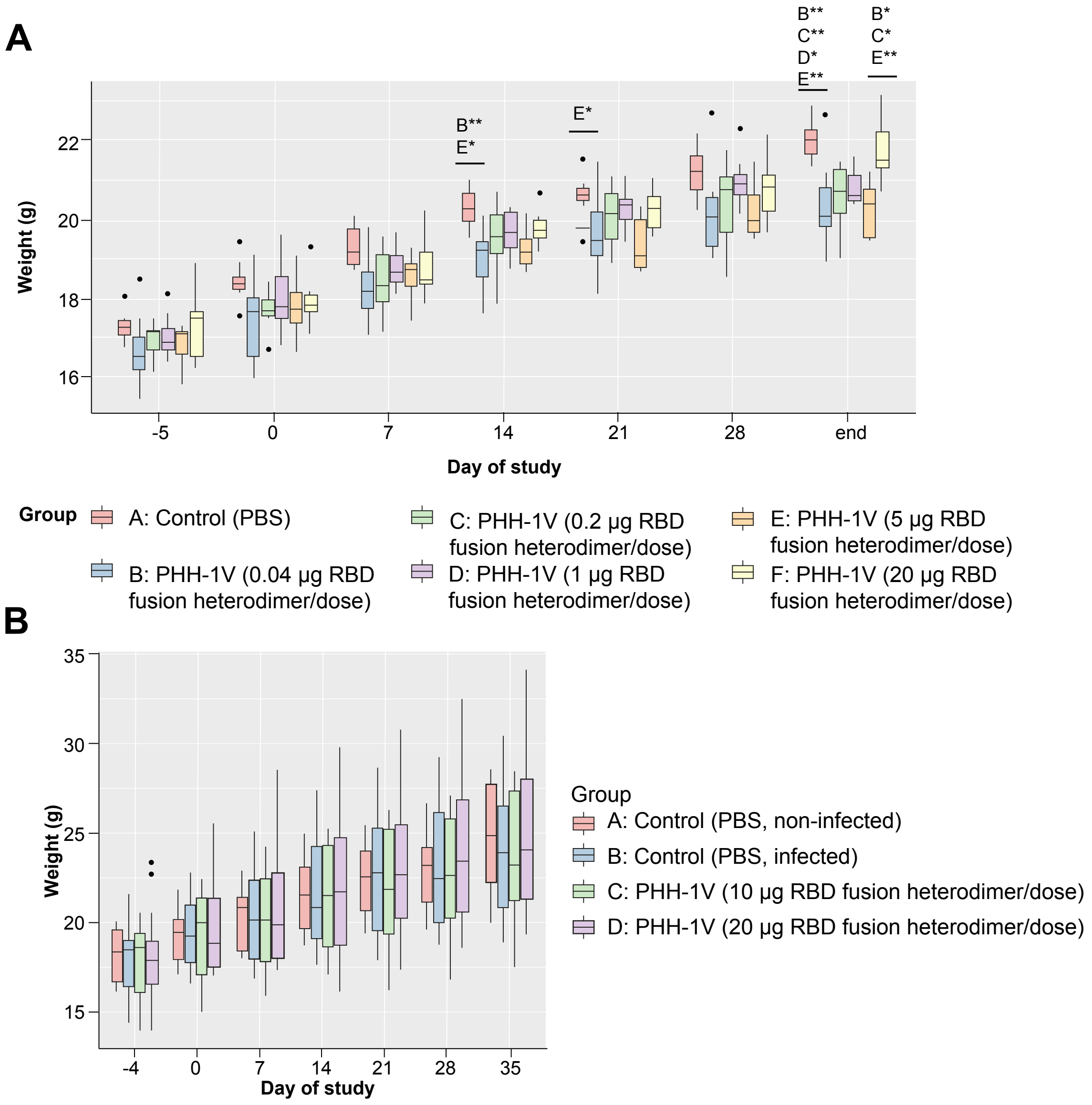


**Figure S1. Bodyweight monitoring during the vaccination period in mice.** Related to Figures 3-5. **(A)** Individual bodyweights in BALB/c mice. Mice were separated into different groups (8 mice/group): group A, vaccinated with phosphate-buffered saline (PBS) as a control group; group B, immunised with the 0.04-µg recombinant protein RBD fusion heterodimer/dose; group C, immunised with the 0.2-µg recombinant protein RBD fusion heterodimer/dose; group D, immunised with the 1-µg recombinant protein RBD fusion heterodimer/dose; group E, immunised with the 5-µg recombinant protein RBD fusion heterodimer/dose; and group F, immunised with the 20-µg recombinant protein RBD fusion heterodimer/dose. Data were analysed by ANOVA from D5 until the end of the study. Data are presented as a box plot: the median marks the midpoint of the data and is represented by the line that divides the box into two parts; the top and bottom limits of the box represent the first and third quartile, respectively; and the upper and lower whiskers represent the lowest and highest values of the distribution, except for the outliers (represented as dots). **(B)** Weekly individual bodyweights of each group during the vaccination period before SARS-CoV-2 infection in K18-hACE2 mice. Group A, vaccinated with PBS and non-infected (n=8, 4F + 4M); group B, vaccinated with PBS and infected with SARS-CoV-2 (n=18, 9F + 9M); group C, vaccinated with 10 µg/dose of recombinant protein RBD fusion heterodimer in oil-based adjuvant and infected with SARS-CoV-2 (n=18, 9F + 9M); and group D, vaccinated with 20 µg/dose of recombinant protein RBD fusion heterodimer in oil-based adjuvant and infected with SARS-CoV-2 (n=18, 9F + 9M). Data are presented as a box plot: the median marks the midpoint of the data and is represented by the line that divides the box into two parts; the top and bottom limits of the box represent the first and third quartile, respectively; and the upper and lower whiskers represent the lowest and highest values of the distribution, except for the outliers (represented as dots).

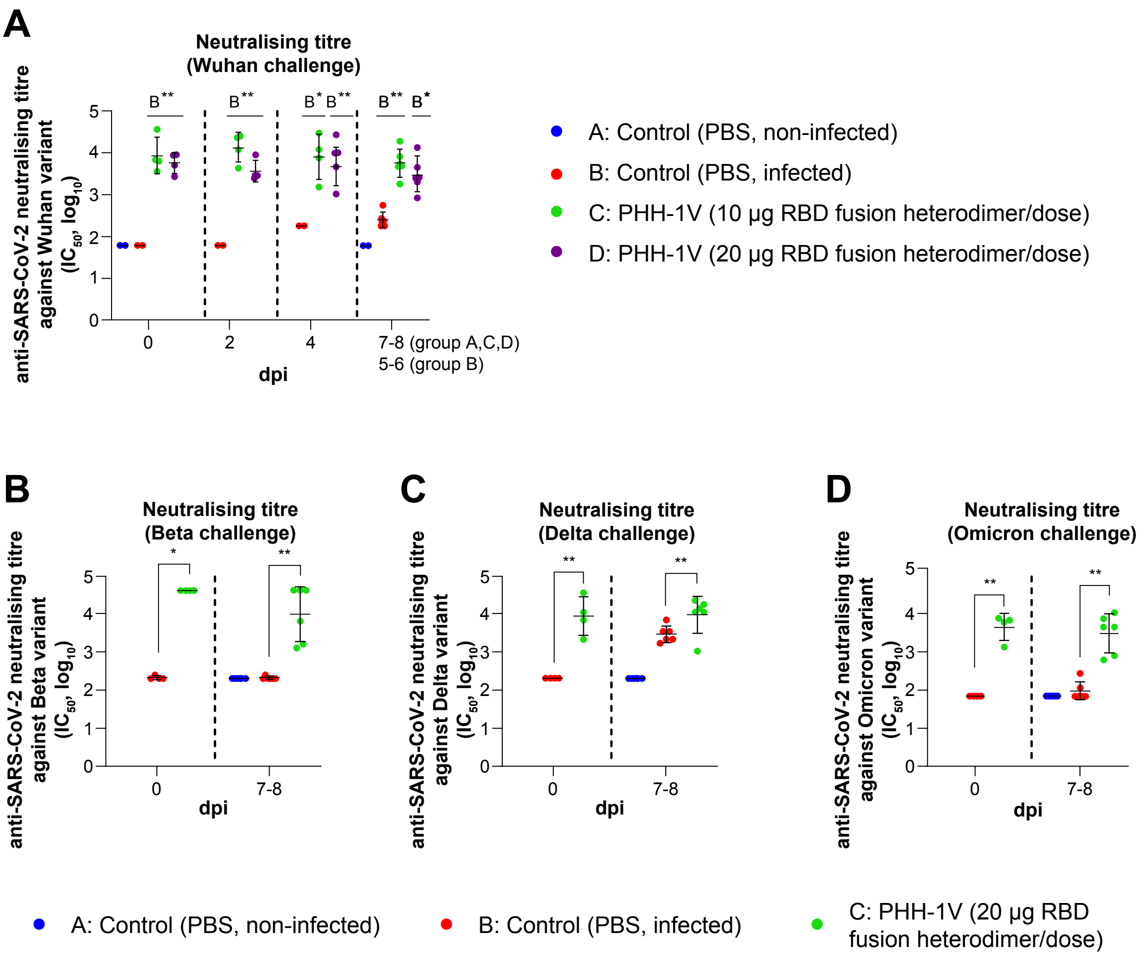


**Figure S2. Neutralising antibody responses by PBNA in K18-hACE2 mice.** Related to Figures 5-8. **(A)** Neutralising titres induced by PHH-1V against D614G Wuhan strain. Group A, PBS-vaccinated non-infected control (n=8); group B, PBS-vaccinated infected control (n=18); group C, vaccinated with 10 µg of recombinant protein RBD fusion heterodimer/dose and infected (n=18); and group D, vaccinated with 20 µg of recombinant protein RBD fusion heterodimer/dose and infected (n=18). Samples of groups A, C and D correspond to 0 (D35), 2 (D37), 4 (D39) and 7 dpi (D42 for males) or 8 dpi (D43 for females); samples of group B were taken 0 (D35), 2 (D37), 4 (D39), and 5 dpi (D40; n=3) or 6 dpi (D41; n=3), when animals reached the endpoint criteria. **(B-D)** Neutralising titres induced by PHH-1V against Beta **(B)**, Delta **(C)** and Omicron BA.1. **(D)** variants. Group A, PBS-vaccinated non-infected control (n=8); group B, PBS-vaccinated infected control (n=18); group C, vaccinated with 20 µg of recombinant protein RBD fusion heterodimer/dose and infected (n=18). All the samples correspond to 0 (D35), 2 (D37), 4 (D39) and 7 dpi (D42 for males) or 8 dpi (D43 for females or at the time of euthanasia in animals reaching endpoint criteria before the scheduled euthanasia day). Titres are expressed as $\log_{10}$ $IC_{50}$. Each data point represents an individual mouse serum, with bars representing the mean titre per group ± SD. These data were analysed using a generalised least squares model on the $\log_{10}$-transformed values of each group. Statistically significant differences between groups are indicated with a line on top of each group: * *p<0.05;* ** *p<0.01*. dpi: days post-infection.

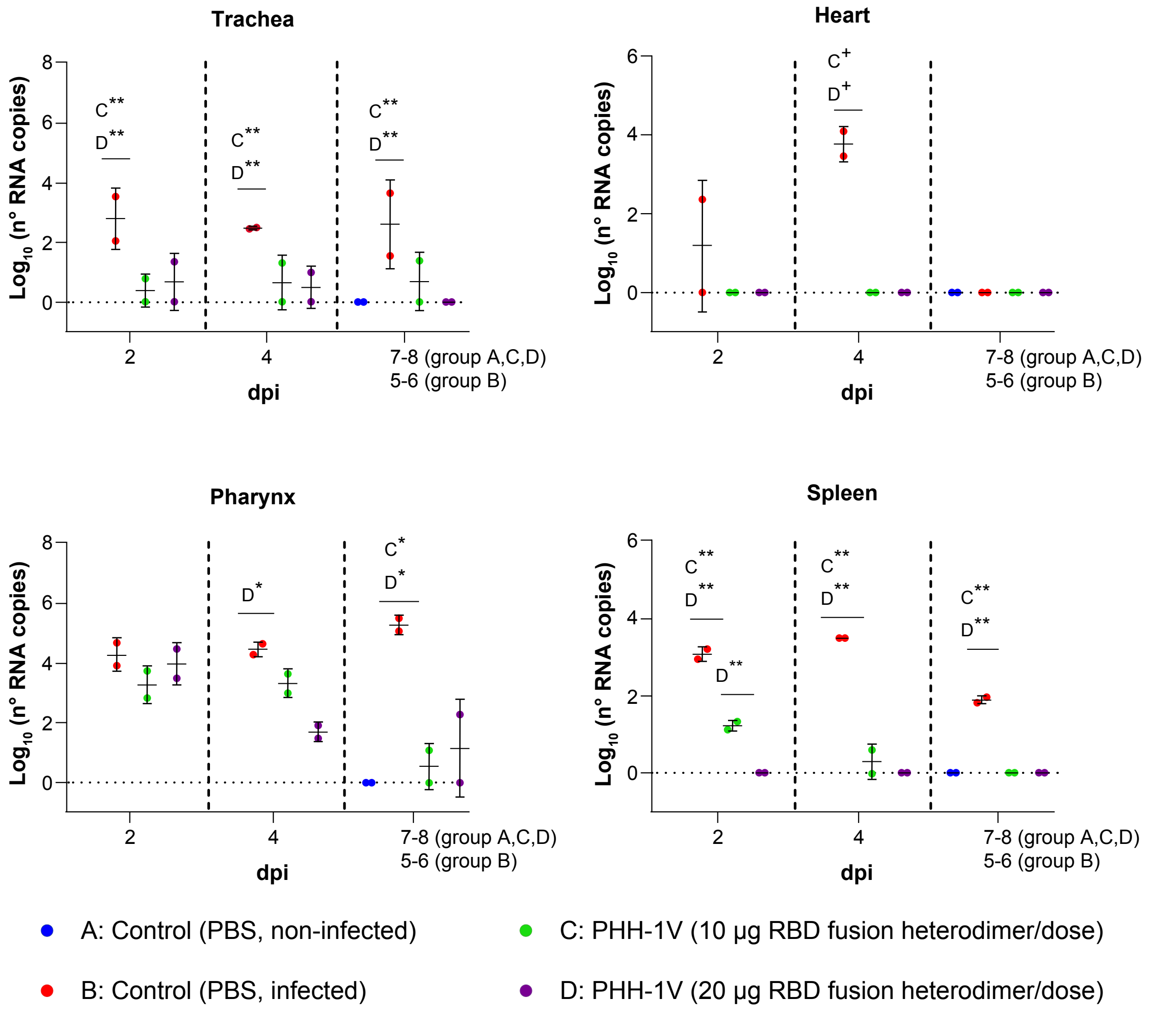


**Figure S3. Viral load in different tissues from mice.** Related to Figure 5. SARS-CoV-2 RT-qPCR detection in trachea, heart, pharynx and spleen in number of copies collected from challenged animals. Each data point represents an individual mouse value, with bars representing the mean ± SD. Samples of groups A, C and D correspond to 2 (D37), 4 (D39) and 7 dpi (D42 for males) or 8 dpi (D43 for females); samples of group B were taken 2 (D37), 4 (D39), and 5 dpi (D40; n=3) or 6 dpi (D41; n=3) when animals reached the endpoint criteria. These data were analysed using generalised least squares models on the $\log_{10}$-transformed values of each group. Comparisons against groups without variability were performed by means of one-sample tests. Statistically significant differences between groups in the number of viral RNA copies are indicated with a line on top of each group: * *p<0.05*; ** *p<0.01;* $^{+}$ *0.05<p<0.1*. dpi: days post-infection.

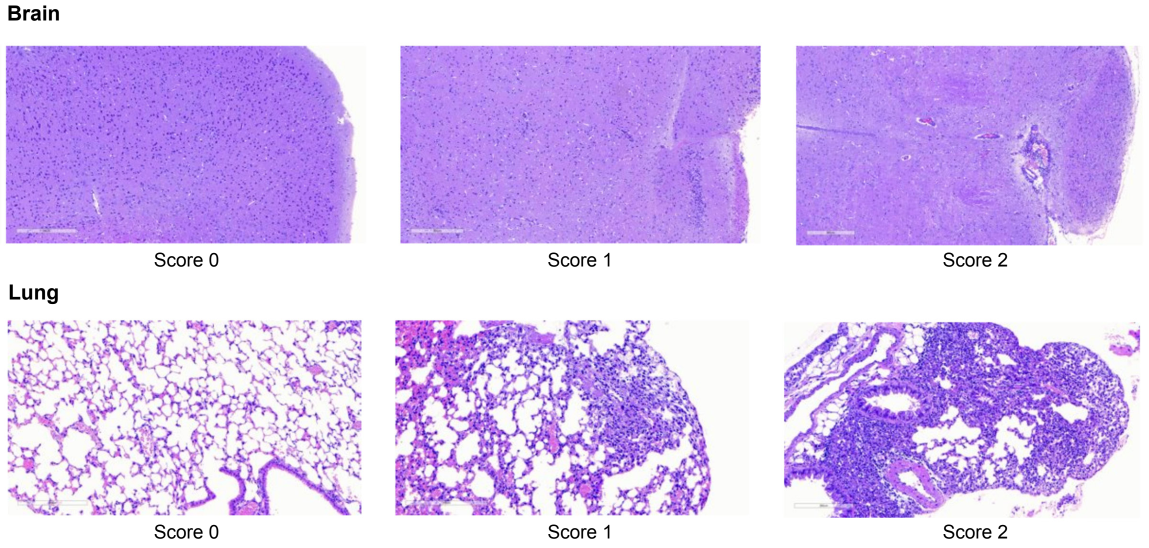


**Figure S4. Representative brain and lung histopathological sections from K18-hACE2 transgenic mice euthanised at days 7-8 post-challenge.** Related to Figure 5. Histopathological section stained with haematoxylin and eosin from brain (top) and lung (bottom) showing scores of 0 (lack of lesions), 1 (mild lesions), and 2 (moderate lesions). None of the study animals showed lesions with score 3 (severe lesions). The brain score section with a score of 0 comes from group A mouse at 7 dpi, the one with a score of 1 comes from group B mouse at 7 dpi, and the one with a score of 2 comes from a group B mouse at 4 dpi. The lung section with a score of 0 comes from group C mouse at 7 dpi, the one with a score of 1 comes from a group D mouse at 7 dpi, and the one with a score of 2 comes from a group B mouse at 4 dpi.

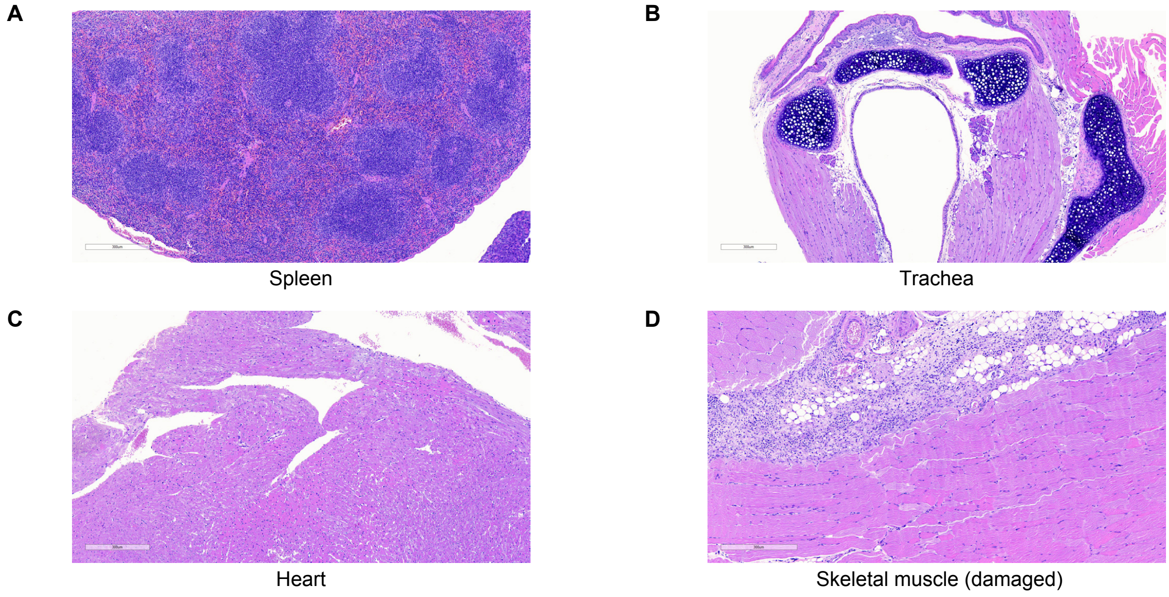


**Figure S5. Representative histopathological sections from K18-hACE2 transgenic mice tissues.** Related to Figure 5. No histopathological lesions were observed in spleen **(A)**, trachea **(B)** and heart **(C)** of SARS-CoV-2 Wuhan/D614G inoculated K18-hACE2 mice at days 7-8 post-challenge. **(D)** Focal mononuclear inflammatory infiltrates in the fascia around muscular fibres of a 20-µg RBD fusion heterodimer/dose on D4 post-challenge with SARS-CoV-2 Wuhan/D614G inoculated K18-hACE2 mice. Haematoxylin & Eosin stain. Bars = 300 micrometres.

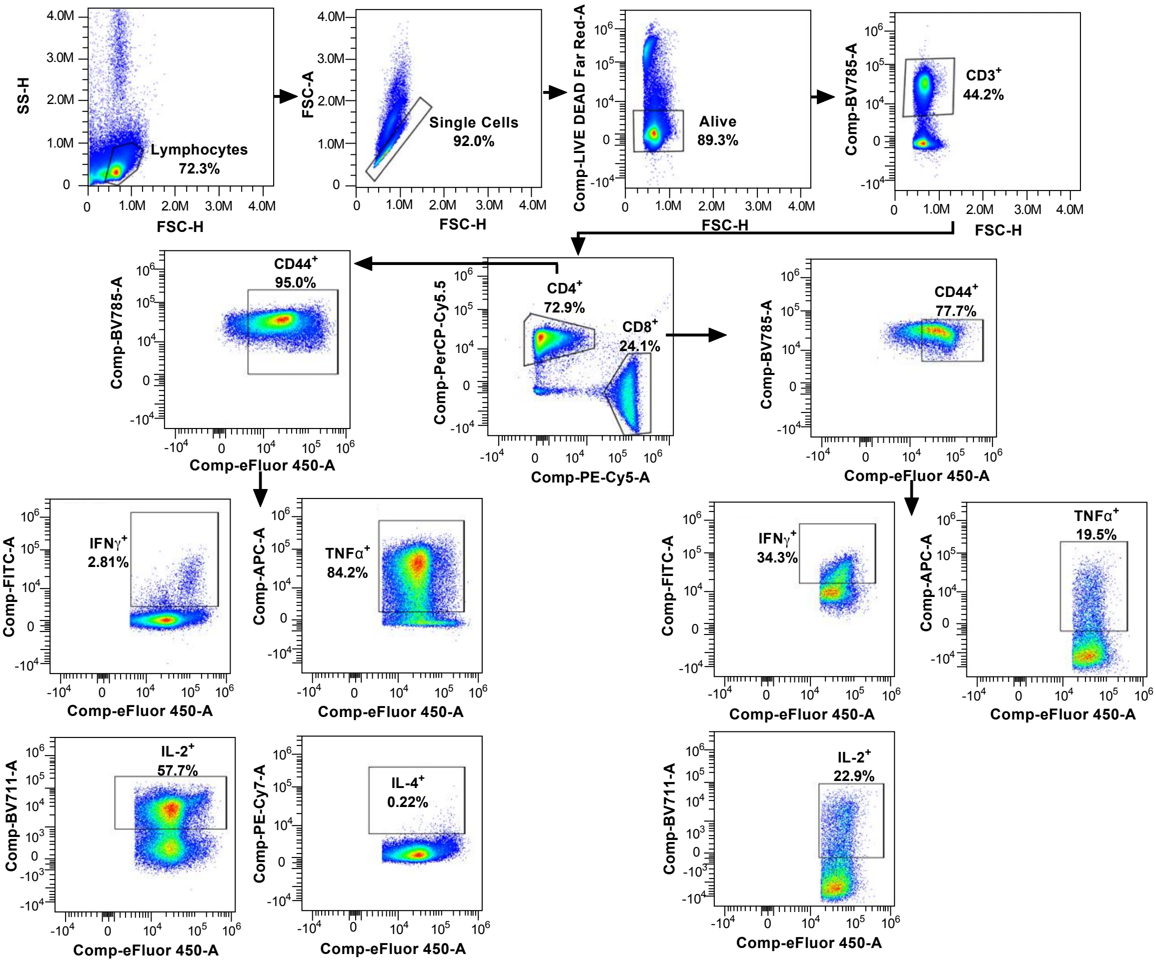


**Figure S6. Identification of IFN-ɣ, TNF-α, IL-2 and IL-4-secreting $CD4^+$ T cells and IFN-ɣ, TNF-α and IL-2-secreting $CD8^+$ T cells from mice splenocytes using flow cytometry.** Related to Figure 4. FSC-H/FSC-A was used to exclude doublets, and T cells ($CD3^+$) within the alive cells (LIVE DEAD Far Red-) were gated. Then IFN-ɣ, TNF-α, IL-2 and IL-4-producing T cells upon mock or RBD-stimulation were measured from $CD4^+CD44^+$ and/or $CD8^+CD44^+$ populations. Fluorescence Minus One (FMO) controls were used to verify flow cytometry data.
